# How well do you know your mutation? Complex effects of genetic background on expressivity, complementation, and ordering of allelic effects

**DOI:** 10.1101/139733

**Authors:** Christopher H. Chandler, Sudarshan Chari, Alycia Kowalski, Lin Choi, David Tack, Michael DeNieu, William Pitchers, Anne Sonnenschein, Leslie Marvin, Kristen Hummel, Christian Marier, Andrew Victory, Cody Porter, Anna Mammel, Julie Holms, Gayatri Sivaratnam, Ian Dworkin

## Abstract

For a given gene, different mutations influence organismal phenotypes to varying degrees. However, the expressivity of these variants not only depends on the DNA lesion associated with the mutation, but also on factors including the genetic background and rearing environment. The degree to which these factors influence related alleles, genes, or pathways similarly, and whether similar developmental mechanisms underlie variation in the expressivity of a single allele across conditions and variation across alleles is poorly understood. Besides their fundamental biological significance, these questions have important implications for the interpretation of functional genetic analyses, for example, if these factors alter the ordering of allelic series or patterns of complementation. We examined the impact of genetic background and rearing environment for a series of mutations spanning the range of phenotypic effects for both the *scalloped* and *vestigial* genes, which influence wing development in *Drosophila melanogaster*. Genetic background and rearing environment influenced the phenotypic outcome of mutations, including intra-genic interactions, particularly for mutations of moderate expressivity. We examined whether cellular correlates (such as cell proliferation during development) of these phenotypic effects matched the observed phenotypic outcome. While cell proliferation decreased with mutations of increasingly severe effects, surprisingly it did not co-vary strongly with the degree of background dependence. We discuss these findings and propose a phenomenological model to aid in understanding the biology of genes, and how this influences our interpretation of allelic effects in genetic analysis.

## Author Summary

Different mutations in a gene, or in genes with related functions, can have effects of varying severity. Studying sets of mutations and analyzing how they interact are essential components of a geneticist's toolkit. However, the effects caused by a mutation depend not only on the mutation itself, but on additional genetic variation throughout an organism's genome and on the environment that organism has experienced. Therefore, identifying how the genomic and environmental context alter the expression of mutations is critical for making reliable inferences about how genes function. Yet studies on this context dependence have largely been limited to single mutations in single genes. We examined how the genomic and environmental context influence the expression of multiple mutations in two related genes affecting the fruit fly wing. Our results show that the genetic and environmental context generally affect the expression of related mutations in similar ways. However, the interactions between two different mutations in a single gene sometimes depended strongly on context. In addition, cell proliferation in the developing wing and adult wing size were not affected by the genetic and environmental context in similar ways in mutant flies, suggesting that variation in cell growth cannot fully explain how mutations affect wings. Overall, our findings show that context can have a big impact on the interpretation of genetic experiments, including how we draw conclusions about gene function and cause-and-effect relationships.

## Introduction

In any given genetic pathway, or even in a single gene, different mutations (whether they are natural variants or lab-induced lesions) can have a wide range of phenotypic effects, and these effects are often modulated by the environment and alleles at other genes throughout the genome (the genetic background) [1–7]. These interactions, however, remain poorly understood. Most studies on genetic background and environmental influences on the phenotypic expression of an allele have examined a single allele of a single gene. As a consequence, a number of questions remain unanswered about how these factors interact to give rise to phenotypic variation, in particular with respect to predictability of those effects [8]. Given our knowledge of context dependence for one allele, can we predict the degree of background dependence for other alleles in that gene or among genes with related functions [9]? Or are the background effects so complex and multifaceted that this remains an impossible task? Moreover, can variation in the phenotypic consequences of multiple alleles, and variable expression of a single allele in different contexts, be explained by similar developmental underpinnings?

Answering these questions is essential for understanding the relationships between genotype and phenotype at a mechanistic level. This also has implications for genetic analysis, *per se*. Many of the genetically based definitions of a “gene” depend on patterns of non-complementation [10]. Likewise, the study of the phenotypic consequences of different alleles of a gene (an allelic series) is necessary for gene structure-function analysis. Investigations of allelic complementation patterns led to discoveries of mechanisms for gene regulation such as pairing-dependence (transvection; [11–13]), position effects [14], and dominant negative interactions [15].

Such genetic inferences—e.g., on gene identity, structure-function relationships, and phenomena such as transvection—rest on the assumption that the phenotypic outcomes of genetic lesions are a function solely of the mutations themselves. However, this view ignores the importance of context dependence, which is relevant when making inferences regarding “higher order” genetic effects. The genetic definition of a gene becomes convoluted if two alleles complement each other only under certain conditions. Likewise, it could be difficult to draw conclusions about the functions of different structural domains in a gene based on a series of alleles with lesions in different locations, if the phenotypic expression of a series of alleles depends on the genetic background or environment. Ignoring the context dependence of allelic effects can influence inferences from mutational analyses designed to dissect the mechanisms connecting genetic lesions to mutant phenotypes.

Nevertheless, the majority of studies examining the influence of genetic background and environment have been limited to a single allele in a given gene. Thus, it is difficult to evaluate how genetic background might influence these higher-level attributes necessary for genetic analysis. Recent studies have highlighted that interactions among mutations can be heavily dependent on the wild-type genetic background in which they are examined [16–21]. Surprisingly, given its central importance in genetics, there has been almost no examination of the influence of wild type genetic background on the ordering of allelic effects or patterns of complementation among mutations within a gene.

There are at least two possible explanations that have been discussed for how genetic background and environment influence penetrance and expressivity; these explanations are not necessarily mutually exclusive and may be viewed as endpoints along a continuum. On one end, context dependence may be unpredictable and highly specific to certain alleles; in other words, knowing how the genetic background and environment influence the expression of one allele does not tell us anything at all about how the same context will influence the penetrance or expressivity of other mutations, even if those mutations affect the same gene or pathway. On the other end, the degree of context dependence is determined by the developmental or physiological constraints of the particular genetic network or trait, not on the unique properties of specific alleles [22]. This perspective on allelic expressivity can be considered an extension of the molecular model for dominance proposed by Kacser & Burns [23], with the robustness being a potentially intrinsic property of a genetic system with thresholds for phenotypic effects. According to this model, the expression of alleles with weak effects (low expressivity) should show little context-dependence, as they minimally perturb the system. Likewise, alleles with very strong effects (e.g., null alleles) should also show little context dependence, because their large effects cause a complete loss of gene function, leaving little room for variability in phenotypic effects due to genetic background or environment. The phenotypic effects of alleles with intermediate effects, on the other hand, are most likely to be sensitive to context. An important corollary to this model is that variability in expressivity and penetrance across genetic backgrounds, and variability within genetic backgrounds, might be correlated; that is, variability across genetic backgrounds and within genetic backgrounds should both be highest for alleles of moderate effect.

Yet despite this reasoning, there is definite evidence for genetic background effects along the spectrum of severity of alleles, including many null alleles [24–27]. At face value, this may suggest that the “intrinisic threshold” model of genetic function may be insufficient to explain genetic background effects. However, this may also reflect that, to our knowledge, no studies have systematically examined an allelic series of mutations that vary along the spectrum of phenotypic effects with respect to genetic background.

Here, we describe a comprehensive, systematic analysis of the effects of genetic background and environmental context on the ordering of allelic series and patterns of complementation using the *Drosophila melanogaster* wing as a model system. We introduced multiple mutations in two genes, *scalloped* and *vestigial*, into two commonly used wild-type genetic backgrounds. Because these two genes interact in a common pathway, this experimental design maximizes the chances of uncovering common influences of genetic background and environment; if genetic background effects are uncorrelated across these two genes, they are not likely to be correlated for other genes, either. We examined how the ordering of allelic effects and patterns of complementation are affected by wild-type genetic background and rearing environment. We demonstrate that the observed genetic background effects are not a property just of specific alleles, but extend across multiple alleles and genes. Genetic background and environmental effects were common, and these influences were most prominent for alleles with intermediate phenotypic effects. However, variability across genetic backgrounds for a given genotype was not strongly correlated with variability within genetic backgrounds. While the relative ordering of allelic effects was consistent across wild type backgrounds, patterns of complementation (intragenic interaction) were consistent only in some instances, differing dramatically in others. Variation in cell proliferation in the developing wing imaginal disc was congruent with phenotypic variation in adult wings among different alleles of each gene. Surprisingly, however, these cellular markers did not always reflect adult phenotypic variation across genetic backgrounds. Our results therefore suggest that multiple distinct mechanisms may underlie variation in expressivity among mutant alleles of the same gene and among genetic backgrounds. We discuss these results both within the broadening context of the biology of the gene, and how to exploit such variation to address fundamental questions in genetics.

## Results

### Despite widespread background and environmental effects on mutant expressivity, the ordering of allelic effects is consistently maintained—

To assess how both wild-type genetic background and rearing environment influenced the expressivity of mutations we used a set of mutations in the *scalloped* and *vestigial* genes (Table 1) that had been repeatedly backcrossed into two common wild type strains, Samarkand (SAM) and Oregon-R (ORE) both marked with the eye color marker *white* (see Materials and Methods). After introgression the strains were genotyped for ~350 anonymous markers throughout the genome including ~100 that distinguished the two wild-type strains. With a few exceptions for particular alleles on particular chromosome arms, introgressions appeared close to complete (Supplementary Table 1, Supplementary Figure 1).

**Table 1.**
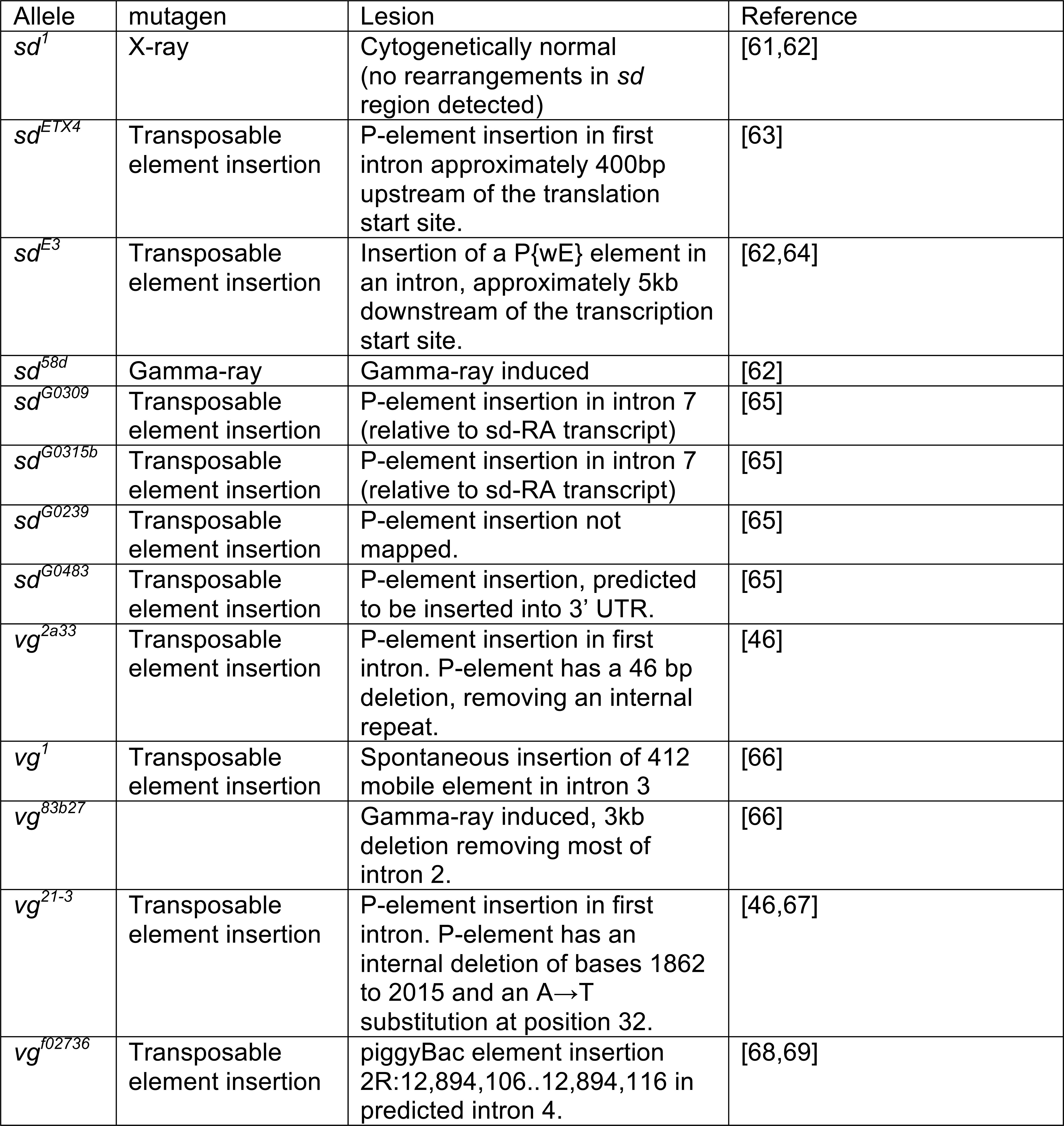
Summary of alleles used in this study.

We observed a striking pattern of genetic background effects related to the overall degree of perturbation caused by the mutations. Some weak and all of the moderate hypomorphic (loss of function) alleles showed genetic background dependence for both genes and at both rearing temperatures. However, the strongest hypomorphs for both *sd* and *vg* showed little or no evidence for sensitivity to the effects of genetic background (Figure 1, 2, Supplementary Figure 2).

**Figure 1.**
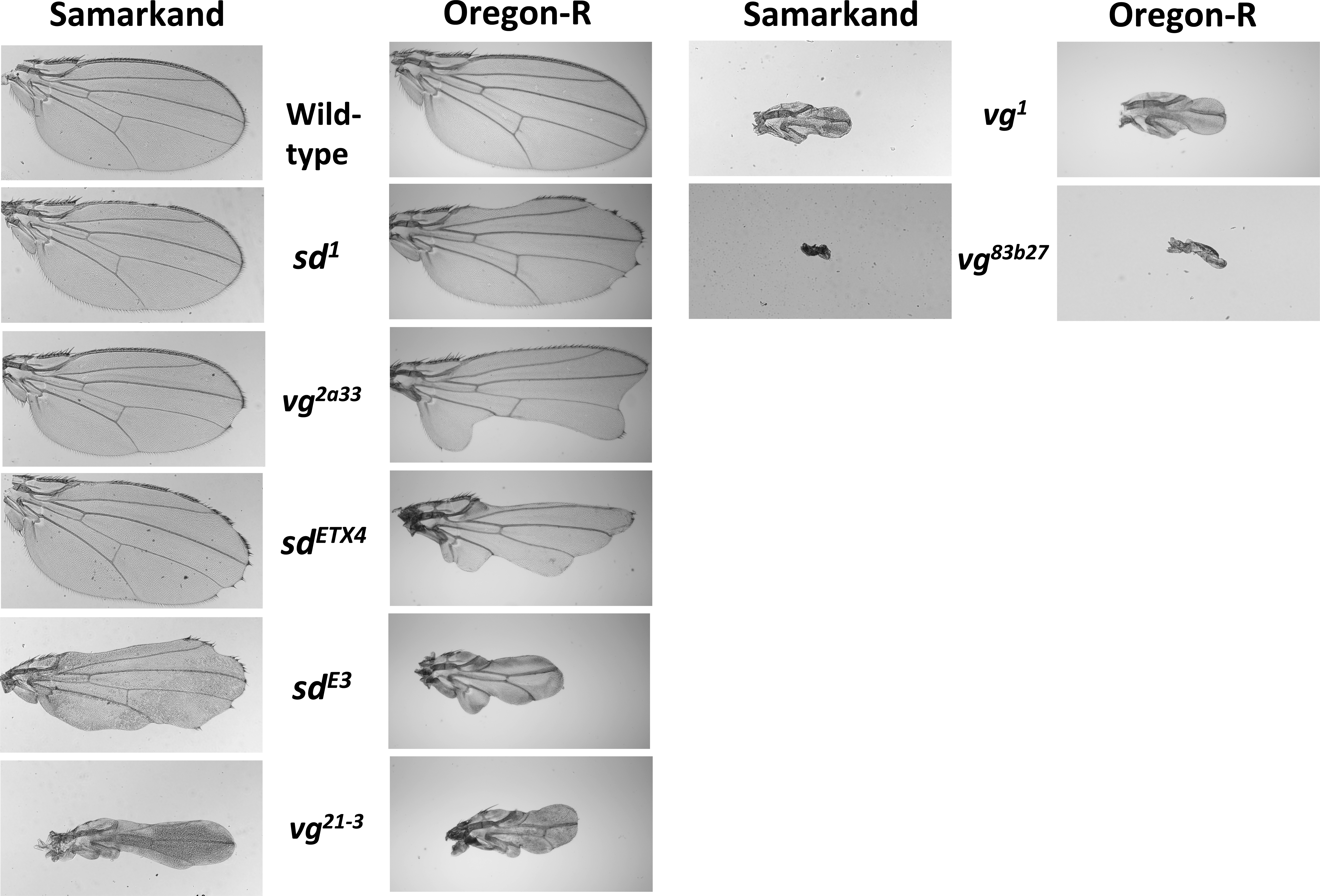
Variation in wing morphology due to allelic and genetic background effects. The left panel shows representative images of wings for a subset of the *vg* and *sd* alleles used in this study that have been introgressed into the SAM wild type genetic background. In the right column are representative images for the same alleles that have been introgressed into the ORE wild type genetic background. While we chose figures to be as representative as possible, it is worth noting that many alleles show a high degree of variability within (as well as between) genetic backgrounds.

**Figure 2.**
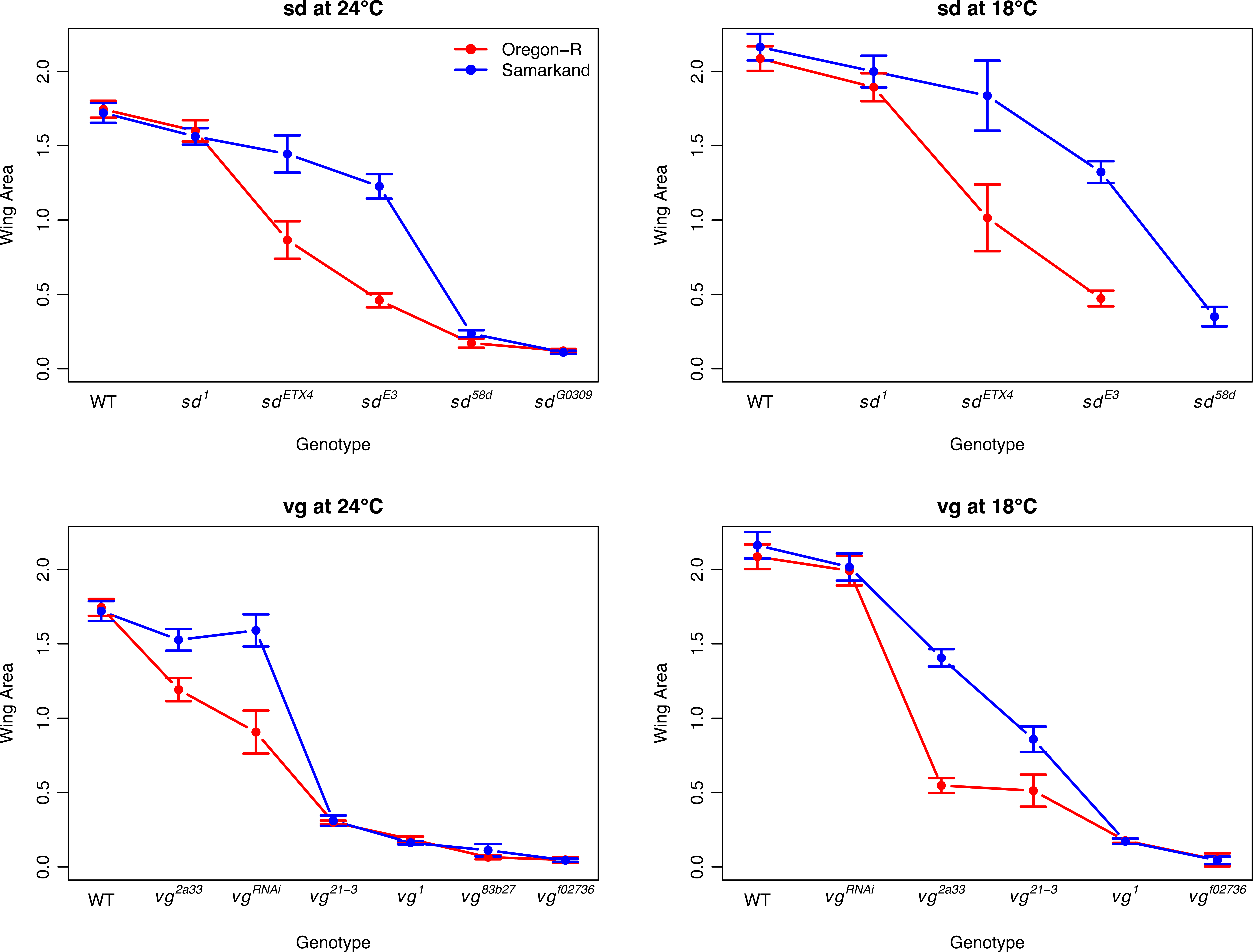
Genetic background and temperature effects of individual alleles of *sd* and *vg* without substantial re-ordering of the rank of allelic effects. The influences of genetic background and rearing temperature on the expressivity of allelic series in the *sd* and *vg* genes, using wing area as a measure of overall wing phenotype. “Stronger” mutations result in smaller wings. The *vg^RNAi^* represents a cross of the UAS_*vg*.RNAi to NP6333-GAL4 (both alleles introgressed into both genetic backgrounds). Error bars represent 95% confidence intervals. See Supplementary Figure 2 for equivalent figures using the semi-quantitative ordinal scale.

Despite the considerable variation in expressivity of the mutations due to genetic background effects, we generally saw consistent ordering of allelic effects. That is, genetic background did not cause substantial changes to the rank ordering of the series of alleles within each gene (Figure 2, Supplementary Figure 2). While there was no overall switching of rank order, certain alleles had largely equivalent phenotypic effects in one genetic background but not in the other, e.g., *sd^1^, sd^ETX4^*, and *sd^E3^*. In some cases, this pattern was seen in some environments but not others (e.g., *vg^2a33^* and *vg^21-3^* show similar effects in Oregon-R but not Samarkand at 18°C, but non-overlapping effects in both backgrounds at 24°C). These results are inconsistent with a model where the wild type genetic background has a constant influence with respect to mutations in the gene. Instead, some non-linear function of both the degree of perturbation and likely the details of the lesion (regulatory vs. coding, etc.) are interacting with genetic background in as yet unexplained ways.

To test whether these results were generalizable to other genetic backgrounds, we crossed the *sd* alleles to 16 randomly selected additional wild type strains that are part of the DGRP collection of sequenced strains [28]. There are a few key differences between this experiment and the more in-depth examination of SAM and ORE. Notably, we did not introgress the mutations; instead, we crossed SAM *sd* mutant females to wild-type males, to obtain F1 flies hemizygous for the *sd* mutation and heterozygous for genetic background alleles on chromosomes 2, 3, and 4. Therefore, the effects of autosomal recessive genetic background modifiers, and all X-linked background modifiers, would not be detected, and we measured background dependence only in hemizygous males. Nevertheless, we observed a similar pattern as before (Figures 1 & 2): while the rank ordering of alleles did not change, some pairs of alleles had similar effects in certain backgrounds and distinct effects in others (Supplementary Figure 3). Thus, this result is likely generalizable across wild-type genetic backgrounds, and depends at least in part on background modifiers with additive and/or dominant effects.

### The influence of background and rearing temperature on patterns of complementation among alleles is complex—

To understand gene structure-function relationships, the analysis of patterns of complementation among alleles of a gene is essential. We adopt the definition of complementation used by [29], in which two mutant alleles complement if, when crossed, they result in phenotypes that quantitatively overlap with wild-type. We investigated how wild type genetic background and rearing temperature influence such patterns (Figures 3 & 4, and Supplementary Figures 4 & 5). The effect of crossing direction on phenotypes was small in magnitude (Supplementary Figures 6 & 7), so for these analyses we treated reciprocal crosses as equivalent. As we observed with the allelic series, the quantitative complementation data suggest that the influence of rearing temperature was relatively modest (Supplementary Figures 4 & 5). In some instances, background dependence has a fairly “constant” effect (same patterns of complementation across backgrounds, but different “intercept”), such as that seen for patterns of complementation between *sd^E3^* and most other *sd* alleles (Figure 3A). Interestingly, this pattern was not exclusively observed. Indeed, we observed some cases where alleles failed to complement each other in one wild type background, but complemented in the other (produced near wild type phenotypes); e.g., at 24°C *sd^ETX4^* and *sd^E3^* complement in Samarkand but not Oregon-R (Figure 3B). Perhaps most interestingly, we observed several cases where hetero-allelic combinations showed background dependent phenotypes, despite lack of background dependence in the homozygotes (Figure 3C); e.g., neither *sd^1^* homozygotes nor *sd^G0309^* homozygotes show much background dependence, but genetic background has a strong influence on phenotypes in *sd^1^*/*sd^G0309^* trans-heterozygotes. In general, we observed that hetero-allelic combinations that resulted in broadly intermediate phenotypic effects were the most background dependent, while those with relatively weak or severe effects had relatively weak background dependence. Thus these results were generally consistent with the observation for homozygous and hemizygous effects of individual alleles for *scalloped* and *vestigial*. Given how generally important the distinction between complementation and non-complementation can be, this pattern seems strikingly and potentially important for inferences in genetic analysis.

**Figure 3.**
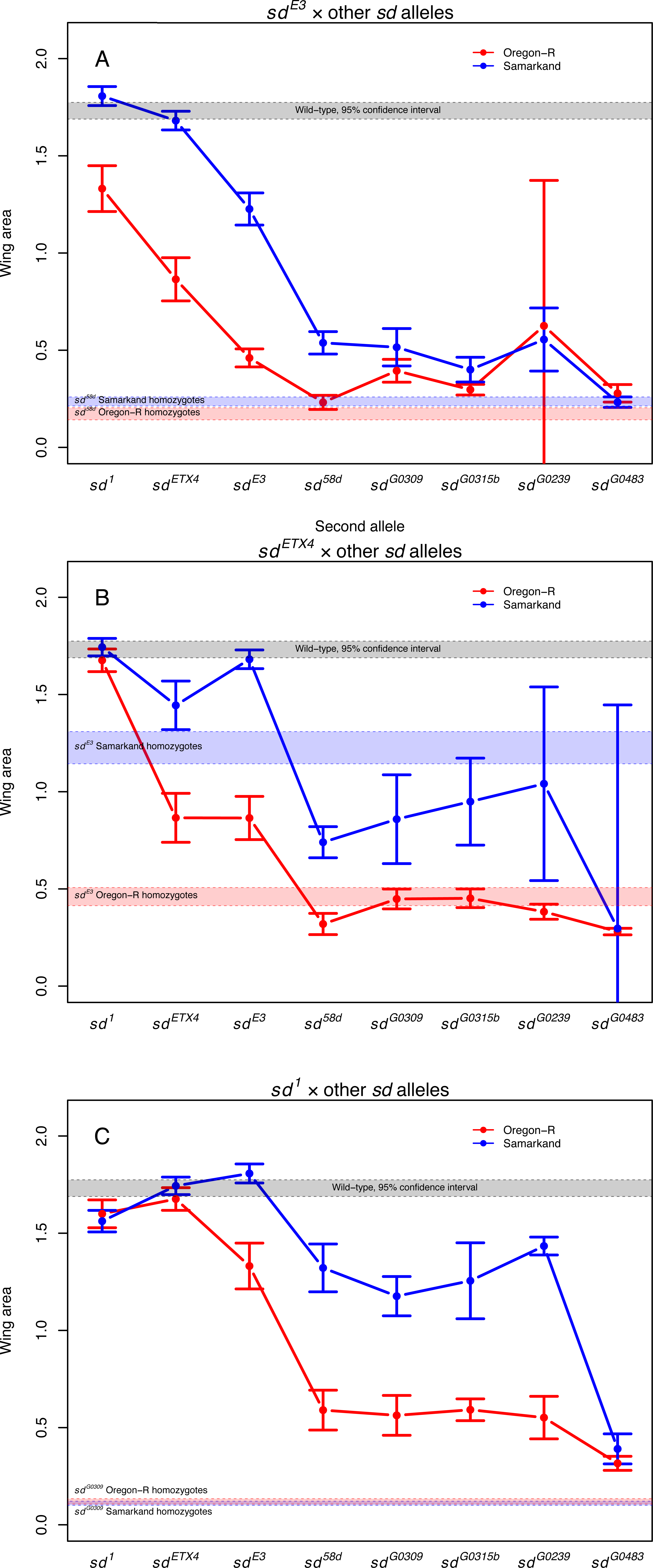
Patterns of intra-genic interactions (complementation) in *sd* at 24°C. Each plot depicts the mean wing phenotype (with 95% confidence intervals) in each genetic background when allele 1 (given at the top of each panel) is paired with allele 2 (x-axis) in diploid flies. (A) Patterns of complementation between *sd^E3^* and other *sd* alleles are largely consistent across genetic backgrounds, although with different “intercepts”. (B) *sd^ETX4^* complements *sd^E3^* in a Samarkand background, but not in an Oregon-R background. (C) Genetic background has little influence on expressivity in *sd^1^* and *sd^G0309^* homozygotes, but a large influence on expressivity in *sd^1^*/*sd^G0309^* heterozygotes. See Supplementary Figure 4 for the complete set of figures for complementation, including ordinal scale measures.

**Figure 4.**
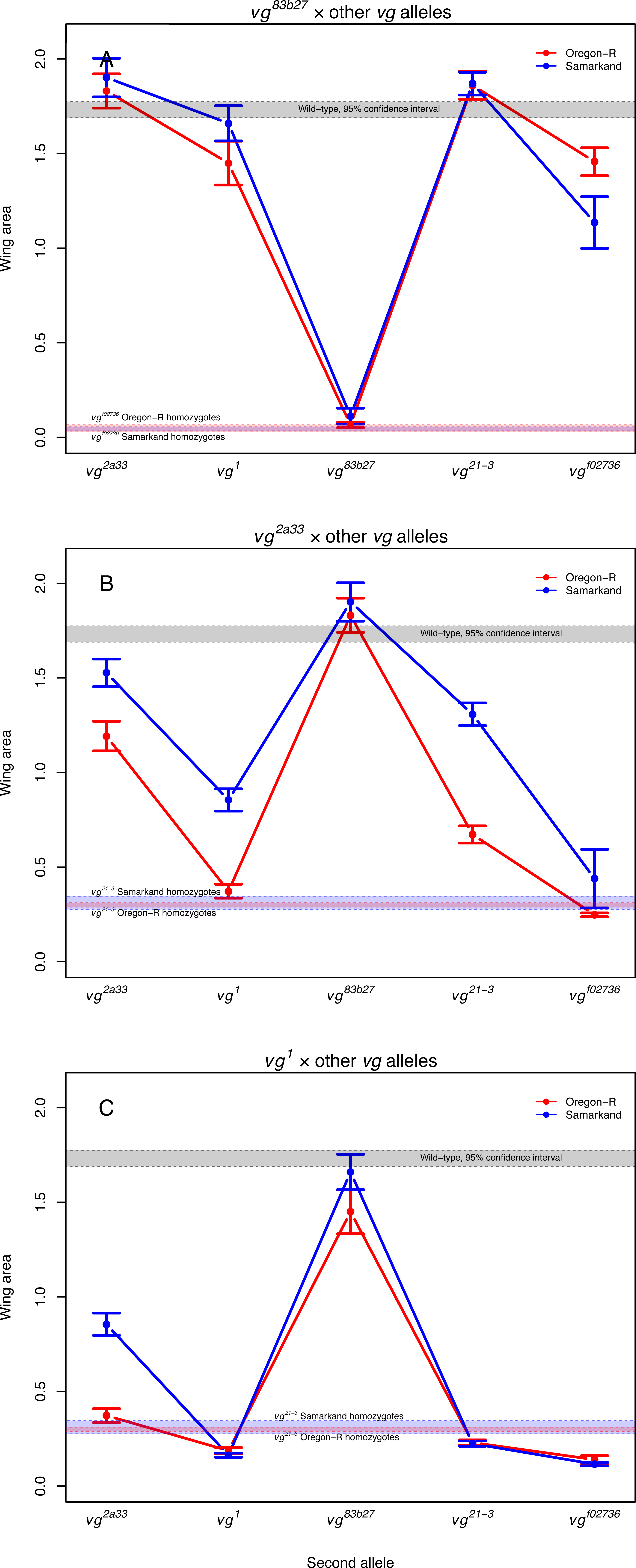
Patterns of intra-genic interactions (complementation) in *vg* at 24°C. Each plot depicts the mean wing phenotype (with 95% confidence intervals) in each genetic background when allele 1 (given at the top of each panel) is paired with allele 2 (x-axis) in diploid flies. (A) *vg^83b27^* exhibits complete or nearly complete complementation with other *vg* alleles, consistent with previously reported patterns of transvection for this allele. (B and C) However, this is not the case for other alleles involved in transvection, such as *vg^2a33^* and *vg^1^*. See Supplementary Figure 5 for the complete set of figures for complementation, including ordinal scale measures.

Two particular alleles of *vg* (*vg^1^* and *vg^83b27^*) have been used extensively to study transvection [30], i.e. pairing-dependent regulation of gene expression as they have mutations in introns 3 and 2 respectively (each providing distinct regulatory sequences). Indeed, among these previous studies there was considerable variation observed in the degree of complementation (compare [3,30] with [13]). Thus, we decided to also examine *vg* allelic combinations including *vg^1^* and *vg^83b27^* for background dependence. As homozygotes both mutations have among the strongest phenotypic effects for viable *vg* alleles (Figure 2, Supplementary Figure 2), yet in combination they have been shown to complement partially or completely [13,31]. We observe largely the same pattern as that reported in the literature (Figures 1 and 2), with surprisingly weak influences of genetic background (Supplementary Figure 2). It also seems as if trans-heterozygotes between other *vg* alleles and *vg^83b27^* also show near complete complementation consistent with this pairing-dependent effects (Figure 4A). This same pattern was not observed in most other hetero-allelic combinations of *vg* (Figure 4B & C, Supplementary Figure 5).

One of these alleles (*vg^1^*) is known to be temperature sensitive; when flies are reared at “low” temperatures (17-20°C) the expressivity of the mutation is strongest, while rearing at higher temperatures (above 25°C) has been reported to result in phenotypes that overlap with wild type [32]. We examined the phenotypic effects of the two most severe (but homozygous viable) alleles of *vg* (*vg^1^* and *vg^83b27^*) in both SAM and ORE reared at three temperatures (18.5, 24 and 28°C). As expected wild type flies (both ORE and SAM) reared at higher temperatures were smaller for wing size (Supplementary Figure 8, squares). However, we observed an intriguing pattern for the *vg1* allele. Flies reared at 18°C and 24°C both demonstrated similar (and severe) phenotypic expressivity with respect to wing size and morphology (Supplementary Figure 8, circles). However *vg1* flies reared at 28°C did show almost completely wild type phenotypes, but only in the SAM background in male flies (Supplementary Figure 8, purple circles). Female SAM *vg^1^* showed a subtle suppression of the phenotypic effects (green circles), while the ORE *vg^1^* showed only a slight (and non-significant) reduction in phenotypic effects (red and blue circles). We also observed weak evidence for temperature sensitivity for the *vg^83b27^* allele (triangles). This suggests that the previously observed temperature sensitivity is not only a function of the *vg^1^* allele, but also depends on how the allele interacts with genetic background and sex.

### Relationship between magnitude of perturbation on wing morphology and background dependence—

As we noted above, the alleles of moderate phenotypic effect seemed to show the strongest degree of background dependence when measured as either homozygotes or hemizygotes (for *sd* males). This suggested a potential relationship between expressivity and sensitivity to conditional effects, or a particular property of the specific alleles. To examine this, we analyzed all of the genetic data (including homozygotes, hemizygotes, and trans-heterozygotes) estimating the variability across wild type genetic background and the average phenotypic effect of the genotype. We observed a pattern resembling an inverted hourglass, where genotypes with weak or severe phenotypic consequences have relatively little background dependence (Figure 5). This includes cases where by themselves the alleles showed no background dependence, but did in combination (Figure 3C), as well as the weakened induction of the *vg*-RNAi (with the NP6333-GAL4) at the lower rearing temperature (Figure 2). This suggests that the pattern is a result of the magnitude of the genotypic effects overall, not a function of specific alleles.

**Figure 5.**
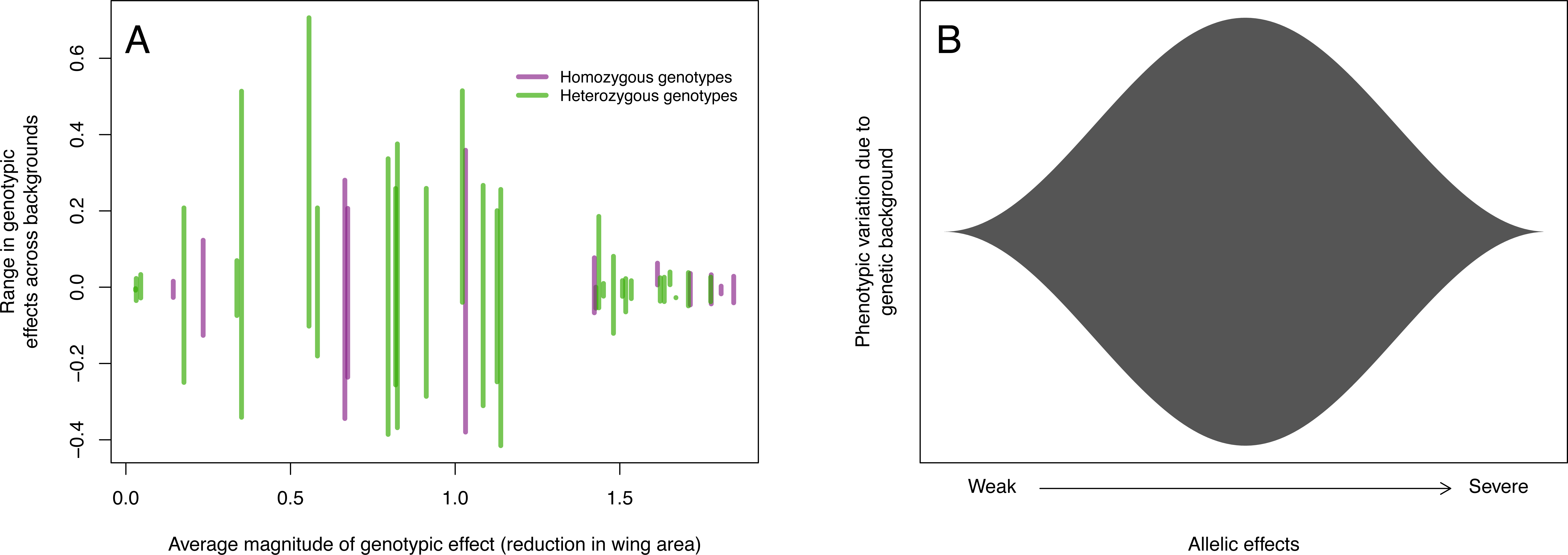
(A) Variation due to genetic background is greatest when mutational perturbation is of moderate phenotypic effect, whereas for mutant genotypes with mild or severe effects on fly wing morphology, genetic background has little influence on the phenotypic effects of the mutation. Each vertical line represents a mutant genotype (e.g., homozygous sd^E3^, or trans-heterozygous vg^1/vg2a33^, etc.). The average effect of a genotype was estimated by computing the difference in wing area (WA) between wild-type flies and flies bearing that genotype (regardless of genetic background, i.e., WA_Mut_ - WA_WT_). The range in genotypic effects across backgrounds was estimated using the difference between the effect in each background and the average effect. (B) Conceptual illustration of the “inverted hourglass” model suggested by this observation. Genetic background effects may be strongest for alleles of moderate effect, whereas genetic background variation is “swamped out” for alleles with very severe effects or for alleles with nearly wild-type function.

Given that we observe variation among genetic backgrounds is greatest for intermediate allelic effects (hemizygous, homozygous or hetero-allelic combinations), we can ask whether this pattern holds with respect to variation within each genotype, i.e. whether phenotypic variability due to micro-environmental variation or developmental noise is also greatest for alleles of intermediate effects. It is well known that variability is higher for many (but not all) mutants relative to their corresponding wild type [22,33]. However it is not clear what relationship is expected for within-genotype variability as a function of the severity of allelic effects. We assessed this by estimating the median form of Levene’s statistic to quantify within genotype and background, but among individual variability [34]. While we did observe a relationship between severity of phenotypic effect and variability, there was no evidence that it was highest for intermediate allelic effects. Indeed the highest within-genotype variability was generally observed for the most severe allelic effects (Supplementary Figure 9B), although there was considerable variation even in such cases. This demonstrates a degree of independence between genetic background effects and the intrinsic sensitivity to micro-environmental variation or developmental noise within genotypes.

### Patterns of cell proliferation are correlated with allelic effects, but do not account for genetic background influences on mutant expressivity—

To assess how the background dependence of these allelic effects are reflected in the underlying developmental processes during wing development, we examined both the size and patterns of proliferation and cell death in wing imaginal discs from mature third instar larvae. Previous work has suggested that mutations in these genes mediate the wing phenotype through their effects on cell proliferation [35,36]. If this is true, we predict that patterns of cell proliferation will vary consistently with severity of both allelic and background effects.

We observed that the Oregon-R wild type wing disc pouches are on average 1.25 times larger than the corresponding Samarkand wild type (Supplementary Figure 10). Yet (and consistent with the observed adult phenotype) the effects of the mutations decrease the size of the pouch far more dramatically (relative to the wild type) in Oregon-R than in Samarkand (Supplementary Figure 10). The effects of *sd* and *vg* alleles on cell proliferation are consistent with previous observations, and with overall magnitude of allelic effects (based on adult wing size). However, cell proliferation in the imaginal disc does not correlate directly with degree of background dependence for adult wing size (compare Figure 2 with Figure 6C,E). Yet despite this, the influence of genetic background is already manifest as shown by the extent of Wingless expression (WG) at the presumptive margin in the wing imaginal disc (Figure 6D,F).

**Figure 6.**
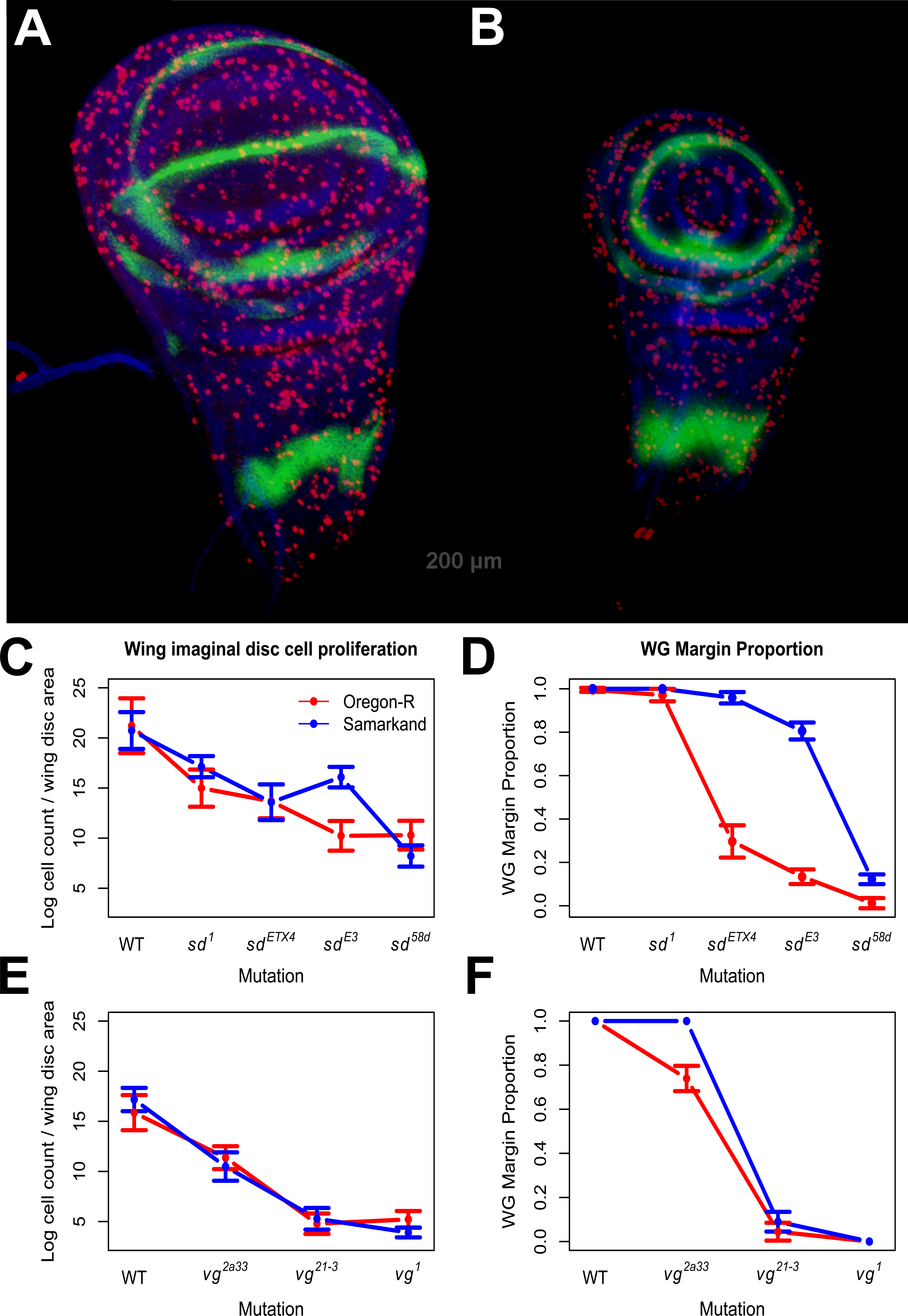
Representative photographs depicting antibody staining for marker of cell proliferation and tissue size for ORE wild type (A) and ORE *vg^1^* (B). Cell number decreases with increasing allelic severity as expected for both *sd* (C) and *vg* (E), but does not reflect differences between genetic backgrounds in mutant expressivity. Thus, while the influence of individual mutations on cell proliferation correlates well with severity of phenotypic effects, it does not match expectations given the background specific consequences of adult wing morphology. However, background dependence is already observed in the wing imaginal disc as demonstrated by the spatial extent of WG expression at the future wing margin for both *sd* (D) and *vg* (F) alleles. Error bars represent 95% confidence intervals.

## Discussion

While both awareness and study of context-dependent allelic effects are becoming increasingly common, most studies have focused on single alleles in a single gene (but see [37,38]). Thus it has remained unclear whether such context dependence is a function of the magnitude of allelic effects, the peculiarities of individual alleles, or the robustness of flux through the genetic network (*sensu* [22,23]). It is also unclear how this context dependence influences other alleles in the gene (and other genes) or patterns of interactions among alleles. We performed a thorough analysis of the contributions of wild-type genetic background and rearing environment on multiple alleles for each of two co-functioning genes. We demonstrate that the background dependence extends to multiple alleles that vary in magnitude of effect and with respect to the nature of the lesion generating the mutant phenotype. Similarly, we demonstrate that the background effects extend beyond the single locus, and can influence alleles at multiple genes that function together (*sd* and *vg*). We systematically examined how genetic background influences the outcomes of higher-order genetic effects, specifically, the ordering of allelic series and patterns of complementation. While phenotypic outcomes were similar across environments and genetic backgrounds in some cases, in others, the genetic and environmental context had strong impacts on results (Figure 2-4). Moreover, variation in expressivity across alleles seems to have a distinct developmental basis from variation in the expressivity of a single allele in different contexts (Figure 6).

One of the most striking patterns requiring explanation is the observation that alleles and genotypes of moderate phenotypic effects showed the greatest sensitivity to genetic background (Figures 2 -5). This was true for homozygous and hemizygous genotypes as well as hetero-allelic combinations. If we had included more alleles of intermediate severity, then we would be likely to observe some rank re-ordering of these moderate effect alleles across backgrounds, despite the overall “anchor” effects of weak and severe perturbations. Some researchers may be tempted to interpret these findings to suggest that mutations of large effect (such as nulls) could be studied with little regard to genetic background effects. However, we would caution against this, depending on the underlying biological explanation for this pattern, of which there are several possibilities. It is unlikely a technical issue of measuring variation for small wings, as we could quantify both adult wing and imaginal disc morphology for all genotypes (e.g., Supplementary Figure 9). Nor is it likely due to the limited genetic variance in the primary experiments (using SAM and ORE) given that we saw similar results across a larger panel of genetic backgrounds (Supplementary Figure 3). One potential source of variation is that introgression is necessarily incomplete (with respect to genomic fragments tightly linked with the focal allele). As such these “ancestral” genomic fragments may also influence the effects with respect to the interaction between the allele and genetic background. However, we think it is unlikely that such effects are having a substantial impact, as the X chromosome does not appear to have any modifiers of the *sd^E3^* background effect [17], and the major region on the second chromosome appears to be independent of *vg* [17,39].

Instead, we favor biological explanations for the “inverted hourglass pattern” that we observed. Such “variational properties” may be intrinsic, resulting from the underlying relationship between the amount of particular developmental signals and the multiple thresholds influencing cell determination and growth, as has been observed in *C. elegans* vulval induction [22,40] and variation under perturbation for otherwise invariant bristles in *Drosophila* [33]. This is analogous to the flux model for the biochemical basis of dominance [23] where there is an underlying threshold for the genetic effect on the phenotype. When gene activity or flux through the pathway is above this threshold (i.e. with very weak mutations) there would be little change in mean phenotype and little context dependence. While severe mutant alleles cause a substantial change in trait mean, they should also create little opportunity for context dependence, as these cause a reduction in gene function or flux to a level sufficiently below the threshold. Thus, allelic or genotypic effects that are of moderate severity near a threshold are mostly likely to be sensitive to context-dependent effects like genetic background. Thus this observed pattern, in which the effects of moderate mutations are more sensitive to genetic background, may be an inherent property of the developmental networks underlying these traits [22,33].

However, there are several observations in both the literature and in the current study that do not fit this interpretation. For instance, previous studies have demonstrated background dependence using null alleles [24], particularly in mice (e.g., [25–27]), though some caution must be used in interpreting “null” effects, as many mammalian genes have paralogs with partial functional redundancy. Additionally, the relationship between sensitivity to genetic background effects and intrinsic variability within a genotype is not strong [41,42]. Indeed in the current study, those alleles with the greatest degree of among-individual, within-genotype variation have the most severe effects on the phenotypic mean (Supplementary Figure 9). Thus the “intrinsic threshold” model described above may be an incomplete explanation for the results observed in the current study.

Alternatively, we speculate there could be an adaptive explanation. Assuming the distribution of mutational effects shows a decrease in frequency with increasing severity, populations will be exposed to weakly deleterious alleles relatively frequently, which could drive the evolution of genetic systems that are robust against weak perturbations. Thus we predict most genetic backgrounds in a population should be able to minimize the phenotypic expression of weakly deleterious mutations. Strongly deleterious mutations occur rarely, and are likely to be purged by selection even when they do occur, so selection for compensatory buffering mechanisms against strong alleles may be ineffective [43]. Therefore, we predict large-effect alleles to have similarly strong effects in most genetic backgrounds in a population. Moderately deleterious alleles, on the other hand, are expected to occur less commonly, so modifier variants that buffer or compensate against these alleles may be more commonly segregating in populations. In other words, some genetic backgrounds in a population will be more buffered than others, so the expression of these moderate-effect alleles may be more dependent on genetic background. However, this speculative model remains to be formally tested.

### Genetic and environmental context influence patterns of allelic interactions—

Collections of alleles are commonly used in experiments designed to dissect gene functions at a fine scale. For instance, using a set of alleles with lesions in different regions allows geneticists to assign functions to specific structural domains or regulatory regions by examining the effects of those mutations on organismal phenotypes (e.g.,[44,45]). Our results suggest that in some cases, repeating such experiments in novel contexts would not necessarily alter overall conclusions, as we did not see any major re-ordering of alleles based upon phenotypic effects. Nevertheless, there were some intriguing differences. For example, some alleles had essentially equivalent effects in one genetic background, but showed differences in the other, and in some cases this difference also depended (albeit weakly) on the rearing temperature (Figure 2). This is interesting given that several *vg* alleles (including *vg^1^*) are known to show patterns of temperature sensitivity [32]. While *vg^1^* did not show changes in expressivity or background sensitivity at rearing temperatures of 18°C or 24°C, we did observe almost wild type-like phenotypes in the SAM *vg1* males reared at 28°C, with much weaker phenotypic suppression for females (Figure 2, Supplementary Figure 8). This suggests that the temperature sensitivity is not a function of just the allele, but due to an interaction between genetic background, sex, and rearing temperature. We also observed weaker phenotypic effects and an increase in background sensitivity in *vg^21-3^* flies when reared at lower temperatures (Figure 2). This allele shows context-dependent phenotypic effects for wing morphology based upon its P-element cytotype [4,46].

While we observed concordance between the effects of *vg* and *sd* across backgrounds, this may not be the norm even for functionally related genes. There is little evidence of correlation across backgrounds for mutational effects of genes involved in the PAR network for lethality associated with early embryogenesis in *C. elegans* [47]; that is, the genetic background modifiers had gene-specific effects, rather than acting similarly on interacting genes. Likewise, the influences of genetic background were incongruent on the phenotypic expression of *sevenless* and *Egfr* mutants in *Drosophila melanogaster* [48], even though these two genes act in related signaling pathways involved in eye development. However, mutations influencing vulval cell induction in *C. elegans* do appear to show moderate concordance across wild type genetic backgrounds [49]. We can think of at least two (non-mutually exclusive explanations) for these contradictory observations. First, as observed in this current study, alleles of different magnitudes can vary in degree of background dependence despite overall concordance between the allelic series in *vg* and *sd*. If previous studies used alleles of large phenotypic effect for some genes, and small-effect for others, then differences in the degree of perturbation may explain the incongruent effects of genetic background, in particular given that perturbation in each gene was evaluated with only one allele or “dose” (for RNAi knockdown). An alternative explanation may be due to subtle aspects of pleiotropy and developmental “degrees of freedom”. In other words, when testing whether the influences of genetic background on expressivity are congruent for different genes, it is important to be sure that the same trait is measured. While the PAR genes discussed above (Paaby et al. 2015) influence the first two cell divisions of embryogenesis in *C. elegans*, the observed lethality may be a result of distinct pleiotropic effects. Similarly, the phenotype scored in the *Drosophila* eye was surface “roughness” (Polaczyk et al 1997), which can be caused by multiple developmental changes, not just transformation of photoreceptor identity [50–53]. This interplay between environmental and genetic context dependence and explaining the degree of phenotypic concordance remains an important area for further research.

As with patterns in the allelic series, while the results of complementation were comparable across contexts in some cases, there were several instances where they differed drastically between genetic backgrounds (Figures 3 & 4). Although certain genetic backgrounds and rearing temperature have similar effects on overall phenotypes across alleles and genes—for example, in this case mutant phenotypes are generally more severe in Oregon-R than in Samarkand—this pattern does not seem to extend to patterns of complementation. Instead, where complementation depends on the genetic background, the outcome seems to involve a complex interaction between the background and the two alleles in question. For instance, *sd^E3^* and *sd^ETX4^* complemented only in Samarkand, and not in Oregon-R. Complementation was observed between *vg^83b27^* and most other *vg* alleles, but not between other pairs of *vg* alleles (similar to [31]), suggesting that transvection is a unique property of this allele or the location of this lesion (*vg83b27* is the only mutation located in intron 2; Table 1). However, we did not observe strong evidence of differences across wild type genetic backgrounds for these allelic combinations, as observed at the *Men* locus [54], although this is likely due to the almost complete complementation observed in combination with *vg^83b27^* (Figure 4). These results have important implications for interpreting genetic analyses, especially if we rely on complementation tests to determine genetic identity, and an important challenge is to determine what causes these differences in complementation. It is important to note that we used a standard definition of complementation (i.e. where the hetero-allelic combination quantitatively overlaps with wild type *sensu* [29]). Arguably complementation could also be defined as any phenotype for the trans-heterozygotes that is quantitatively more like wild type than the homozygotes (i.e. over-dominance). However, this definition does not substantially alter our conclusions.

### Correlation between expressivity and cellular phenotypes for allelic series and background effects, and interpreting causal links—

We examined several developmental and cellular correlates of the phenotypic effects: patterns of cell proliferation, overall size of the developing wing, and development of the future margin in the mature third instar wing imaginal discs. Previous work has described the patterns of reduced cell proliferation [35,36,55] associated with mutations in the *sd* and *vg* genes. Indeed the associations between the effects in the wing disc and the morphology of the adult wing for these mutations may suggest a causal relationship, as viewed across the allelic series in either background. However, the results presented here caution against making such causal inferences too quickly (Figure 6). A clear relationship exists across alleles between the severity of the adult phenotype and the amount of proliferation in the imaginal disc (Figure 6C,E). Yet it is clear that the strong background dependence of the observed effects on adult morphology is not reflected in this cellular correlate during development. For example, both *sd^ETX4^* and *vg^2a33^* genotypes have much smaller wings in Oregon-R compared with the Samarkand background (Figures 1 & 2). Yet cell proliferation in the wing disc is indistinguishable between backgrounds for these genotypes (Figure 6C,E). In contrast, differences among the genetic background are observed using WG expression in the imaginal disc (concurrent with the marker for cell proliferation), showing that genetic background effects are already present (Figure 6D,F). Thus variation in proliferation accounts for only part of the effects observed for adult wing morphology. This is reminiscent of earlier work with mutations in the *mushroom body miniature* gene, which showed that a mutation's effects on the size of the mushroom bodies (part of the insect brain) and its effects on learning were in fact separable across two different wild type genetic backgrounds [56]. Indeed, studies of mutational effects across wild-type genetic backgrounds may be a useful tool to distinguish true causality from so-called epistatic pleiotropy [57].

### A plea for maintaining “legacy genetics toolkits”—

With the advent of tools using RNAi, over-expression, full gene knockouts, targeted deletions and direct allelic replacements, the experimental capabilities of most geneticists have substantially expanded in recent years. In comparison the “legacy” mutational toolkits that were available for the first 80+ years of genetics research analysis were generated in a less standardized fashion, with uncertainty remaining for many alleles with respect to many aspects of function. Given the costs associated with maintaining such collections it may seem like allelic variants (such as those used in the current study) may not be important to maintain. Nevertheless, if the newer toolkits are used at the exclusion of “legacy” mutations, the new tools may ironically lead to a decrease in the diversity of genetic research. Without many of the baroque facets of mutational approaches that have been employed in the past, it is possible that phenomena like transvection and position effect variegation may have never been discovered. These in turn have had a profound influence on our understanding of gene regulation.

### Material and Methods

#### Fly strains and Introgression of alleles —

The two wild-type strains used for this study were Oregon-R (ORE), and Samarkand (SAM), both marked with a *white* (*w*) allele and are maintained as inbred lines, and are regularly genotyped to avoid any contamination, and maintain homozygosity [17,19,58]. The origin of mutant alleles used in this study can be found in Table 1. Introgression of the alleles was performed largely as previously described for other alleles [17,19]. However, in several instances, balancer chromosomes (that had been previously introgressed into each wild-type background) were used to transfer entire chromosomes, followed by backcrossing to complete introgression. After completion of introgression all strains were genotyped for ~400 markers, of which ~100 explicitly distinguish the two wild-type strains used in this study (Supplementary Table 1, Supplementary Figure 1). While most alleles showed upwards of 90% introgression, some alleles still showed some degree of the ancestral background. Thus, we recognize that in additional to the focal alleles, there are some small genomic regions from the ancestral backgrounds that will contribute to the observed effects. We argue that this final step (genotyping) should be employed whenever possible, as there is no guarantee of successful introgression of the majority of the genome, and residual genomic fragments from the progenitor background may result in artifacts.

While almost all alleles used in this study represent “classic” alleles (in that the effect is a result of mutation at the native locus), for vg we were concerned about having insufficient phenotypic coverage (in terms of severity). Thus we introgressed a UAS-*vg*.RNAi (VDRC ID 16896) and NP6333 (Pen)-GAL4 into both Samarkand and Oregon-R. However, it is worth noting that the temperature dependent effects of this UAS-GAL4 combination likely reflect the temperature sensitivity of GAL4 *per se*.

To determine whether the patterns observed across the Samarkand and Oregon-R wild type backgrounds could be generalized, we crossed three of the *sd* alleles (*sd^1^, sd^E3^* & *sd^58d^*) as well as the corresponding SAM wild type to 16 of the sequenced wild type lines that are part of the DGRP collection [28]. As the original SAM *sd^58d^* was lost between the first and subsequent experiment, this mutations was re-introgressed independently (starting from the ORE *sd^58d^*). After introgression, the phenotypic values were compared and found to be indistinguishable from the original estimates of SAM *sd^58d^*.

#### Experimental crosses—

All mutant and wild-type flies in Oregon-R as well as Samarkand genetic background were separately raised at low density, in bottles with ~50ml fly food at both 24°C and 18°C with 65% relative humidity and 12 hr light/dark cycle (Percival, Model: I41VLC8, incubator). After maintaining the fly strains in these conditions for at least 2 generations, virgin females and males were collected and housed separately in vials with 40 individuals/ sex/ genotype/ background at 24°C and 18°C respectively. For every strain, this collection was performed for a total period of 4 days upon eclosion, under CO2 anesthesia and the flies were maintained in vials (as above) for an additional 3 days to remove any residual effects of the CO_2_. Following this, all the flies in a given treatment were randomized (within genotype) and 40 pairs were then allowed to lay eggs on 35mm x 10mm cell culture plates with grape juice agar (2% agar in 50% grape juice:water) for 15-22 hours. For generating the allelic series data, 40 pairs of flies per allele were crossed among themselves and to generate the intragenic complementation data, 40 pairs of chosen genotypes were crossed to each other. Eggs collection was performed from multiple such plates per treatment and 4 replicates were created with each replicate having 40 eggs in a vial with ~10-12ml fly food/ cross/ background at both 24°C and 18°C respectively. These eggs were allowed to develop at the respective temperatures, and upon eclosion the adults were stored in 70% ethanol for further analyses. However, we have no data for some specific genotypes, as they were lethal in one or more combinations of background and rearing temperature. All of these experimental crosses were performed simultaneously using a single batch of media. While for the vast majority of experimental crosses our design provided sufficient samples (i.e. at least 10 individuals per genotype/replicate vial), several alleles showed partial lethality, and thus for some alleles and allelic combinations we were only able to collect a small number (~3-5) of individuals. For experimental crosses that did fail to yield F1 progeny during the primary experiment, an additional block of crosses (Block 3) was setup. As controls for block level effects, wild type and a few (randomly chosen) mutant crosses that were successful in the initial experiment were also setup in this block. For this additional setup, wild type and relevant mutant flies in both backgrounds were grown, and virgin females and males were subsequently collected as described above. Four replicate vials, each containing six virgin females and four males of the appropriate genotypes were created. The flies were allowed to mate and lay eggs for 3-4 days following which the adults were discarded and the vials were allowed to develop and were further processed as described above.

For crosses of the DGRP strains to the *sd* mutations introgressed into Samarkand (and the corresponding Samarkand control) we crossed females bearing the *sd* alleles to males of each of the 16 DGRP lines and reared these in the incubator described above at 24°C. When the F1 progeny emerged the male F1 *sd* hemizygotes (*sd* is X-linked) were stored in 70% ethanol for phenotyping.

To further investigate the joint effects of rearing temperature and genetic background on the phenotypic expressivity of the *vg1* and *vg^83b27^* alleles we reared the strains bearing these mutations in both Oregon-R and Samarkand (as well as the corresponding wild types) at 18°C, 24°C and 28°C. For the 28°C treatment a Fisher Scientific incubator (Model 3070C) was used, but with otherwise similar settings to the Percival incubators.

#### Wing Imaging and quantification—

A single wing was dissected from at least 5 individuals/sex/genotype for 2 replicates and mounted in 70% glycerol for a total of at least 10 observations/ genotype. Images of the wings were captured using an Olympus DP30BW camera mounted on an Olympus BW51 microscope using DP controller image capture software (v3.1.1). The wing area was then obtained using a custom macro in ImageJ software (v1.43u). Measures of wing area can be confounded by variation for body size. Furthermore, some mutations have subtle effects only causing bristle loss at the wing margin, but with no influence on wing size. Both of these issues are of particular concern for weak hypomorphic alleles (Figure 1). Therefore, in addition to using wing area, we also utilized an ordinal scale to measure severity of the phenotypic effect (on a scale of 1-10). This approach [19,59] has been used successfully and correlates well with wing area.

For the crosses to the DGRP lineages, wings were imaged using a Leica M125 Microscope (50X total magnification) and captured with a Leica DFC400 Camera. For the additional crosses of the *vg^1^* and *vg^83b27^* alleles the images were captured on an Olympus microscope with an Olympus DP80 digital camera (40X total magnification). For all estimates we generated genotypic means and 95% confidence intervals. To examine the amount of variation within genotypes, we used two forms of Levene’s Statistic, using the genotypic medians, and then use the absolute value of the differences (or log transformed values of these values). See [34] for further discussion on these methods. We did not rescale the among background effects for the primary experiment (i.e. using CV) given some of the known assumptions that are violated (i.e. potential lack of a linear relationship between standard deviation and the mean under perturbation for a trait). See [22,60] for more details. In particular for the main experiments, because we had only two genetic backgrounds, estimates of the among background variances would be poor. However, for the subsequent follow up experiment with the 16 DGRP strains we did estimate the between-background variability using Levene’s statistic as described above.

#### Immuno-histochemistry of wing imaginal discs—

10-15 larval heads with the wing imaginal attached were dissected from wandering third instar larvae in 1x Phosphate Buffered Saline (PBS) per genotype. These were then fixed in 4% paraformaldehyde dissolved in 1x PBS for 20 minutes at room temperature. This was followed by repeated gentle washing (4x) for 15 minutes each with a PBT solution consisting of 1x PBS and 0.1% Tritonx100 at room temperature. After washing, the tissues were treated with a PBTBS blocking solution consisting of PBT, 0.1% Bovine serum albumin and 2% Goat serum for 2–3 hours at 4°C. This was then followed by overnight (minimum of 4 hours) incubation with the primary antibody at 4°C on a shaker followed by washing as above. This was followed by blocking (2 hours), overnight (minimum of 4 hours) incubation with secondary antibody (at 4°C) and washing as above. Post-washing, the tissues were incubated with Hoechst stain (Sigma) at 1:15000 dilution (blue color), for ~1 hour, the discs separated and mounted on a slide. Images were captured using Olympus DP30BW camera mounted on an Olympus BW51 microscope using the DP controller image capture software (v3.1.1). The images were merged using the ImageJ software (v1.43u). The primary antibodies used were mouse anti-WG (4D4, 1:500 dilution) or anti-ct (2B10, 1:100) from the Developmental Studies Hybridoma Bank used in conjunction with rabbit phospho-histone H3 (1:500, Cell Signalling Technology). Secondary antibodies were anti-mouse FITC (1:1000) and anti-rabbit DyLight 594 (1:500). At least 10 wing discs were imaged for each genotype x background combination (1 image per channel) on the same microscope described above. We then applied custom ImageJ macros to the image stacks to estimate the area of the wing pouch (defined by the proximal ring of WG expression), and count phospho-histone H3 positive cells in that region. Since *sd* and *vg* influence the development of the wing margin, we measured the length of the central part of the imaginal disc as defined by the central WG expression band (the future wing margin) by using the line tool in ImageJ. In cases where the bands were incomplete or contained gaps, a sum of lengths of all the individual bands was used and for discs that lacked the central band, we measured the length of the central part. We then determined the proportion of wing margin length to the length of the complete margin (the curve from anterior to posterior) for each of the *sd* and *vg* alleles. Refer to figure 6A to see a complete WG expressing margin.

## Acknowledgements

We would like to acknowledge all of the fly labs who provided alleles for this study, without which this work would have never been possible. Thanks to Lindy Johnson and Heather McGovern for help with fly work. We would also like to thank Dr. Trudy Mackay, Dr. Gregory Copenhaver and three anonymous reviewers for valuable feedback that has improved this manuscript.

---

**Supplemental Table 1.**
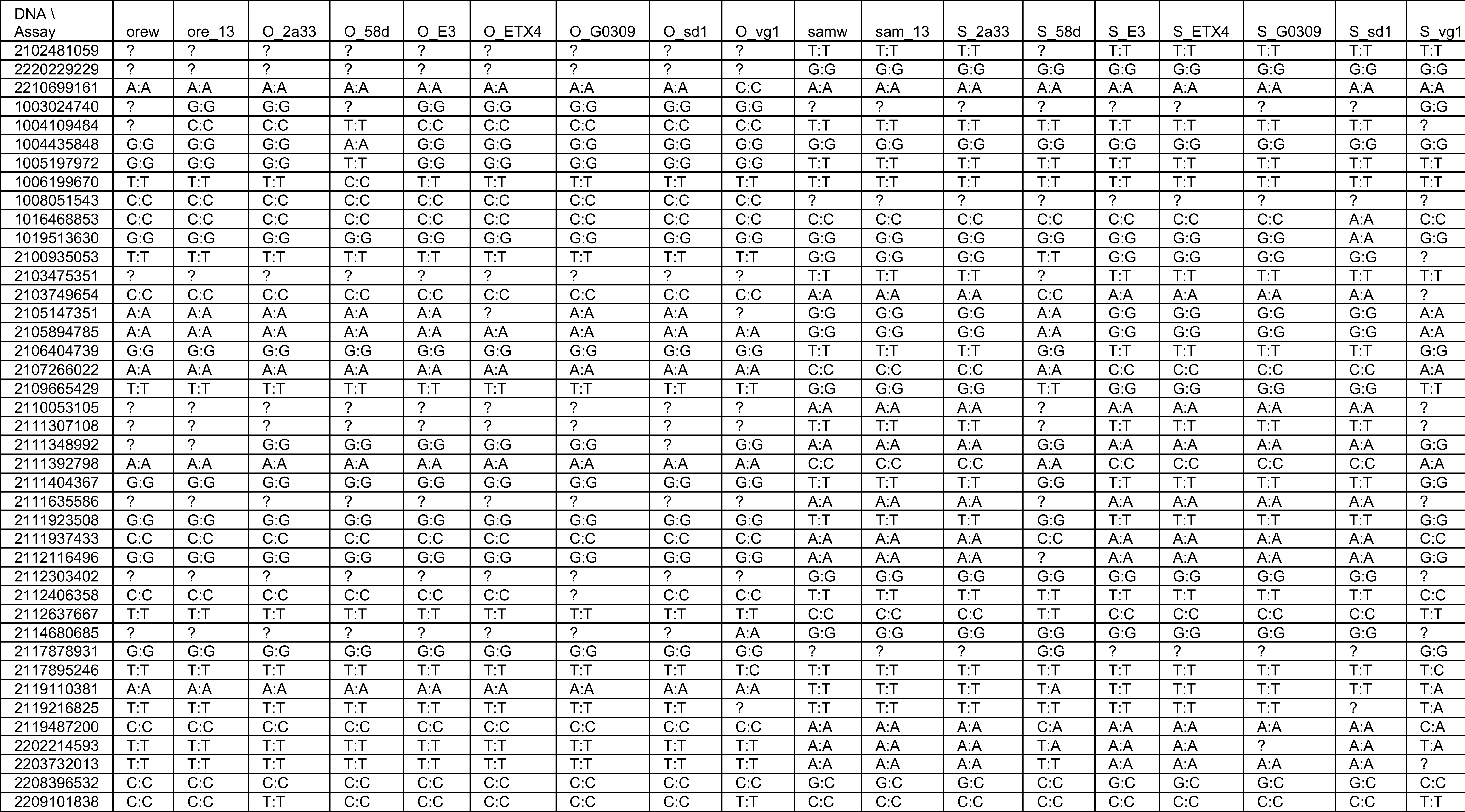
Marker genotyping showing introgression of mutant alleles into 932 o the 933 Oregon-R and Samarkand wild genetic backgrounds is nearly complete.

**Supplemental Figure 1.**
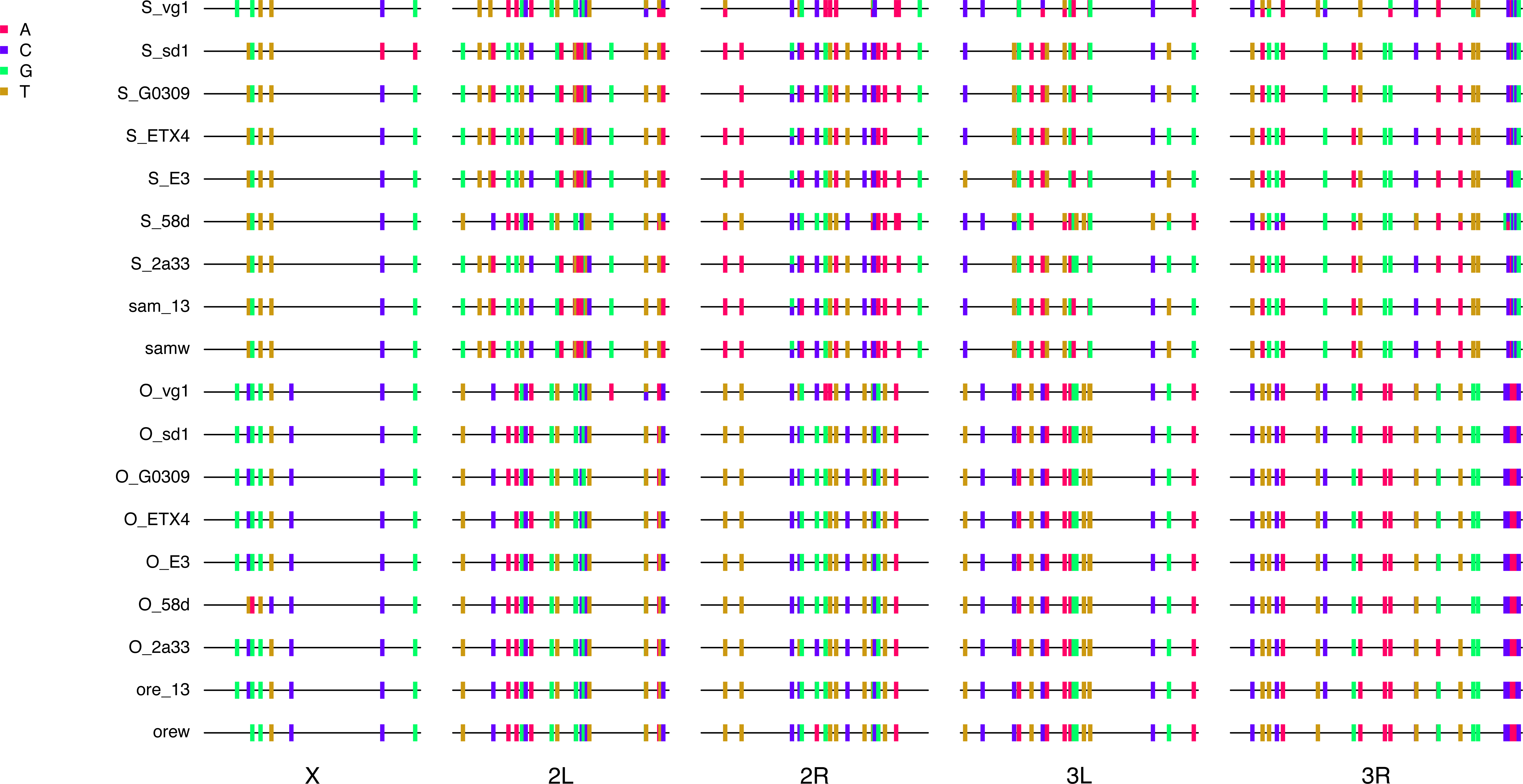
Genotypic assessment of introgressions. With a few exceptions, 936 introgressions appeared mostly complete, except close to the focal allele.

**Supplemental Figure 2.**
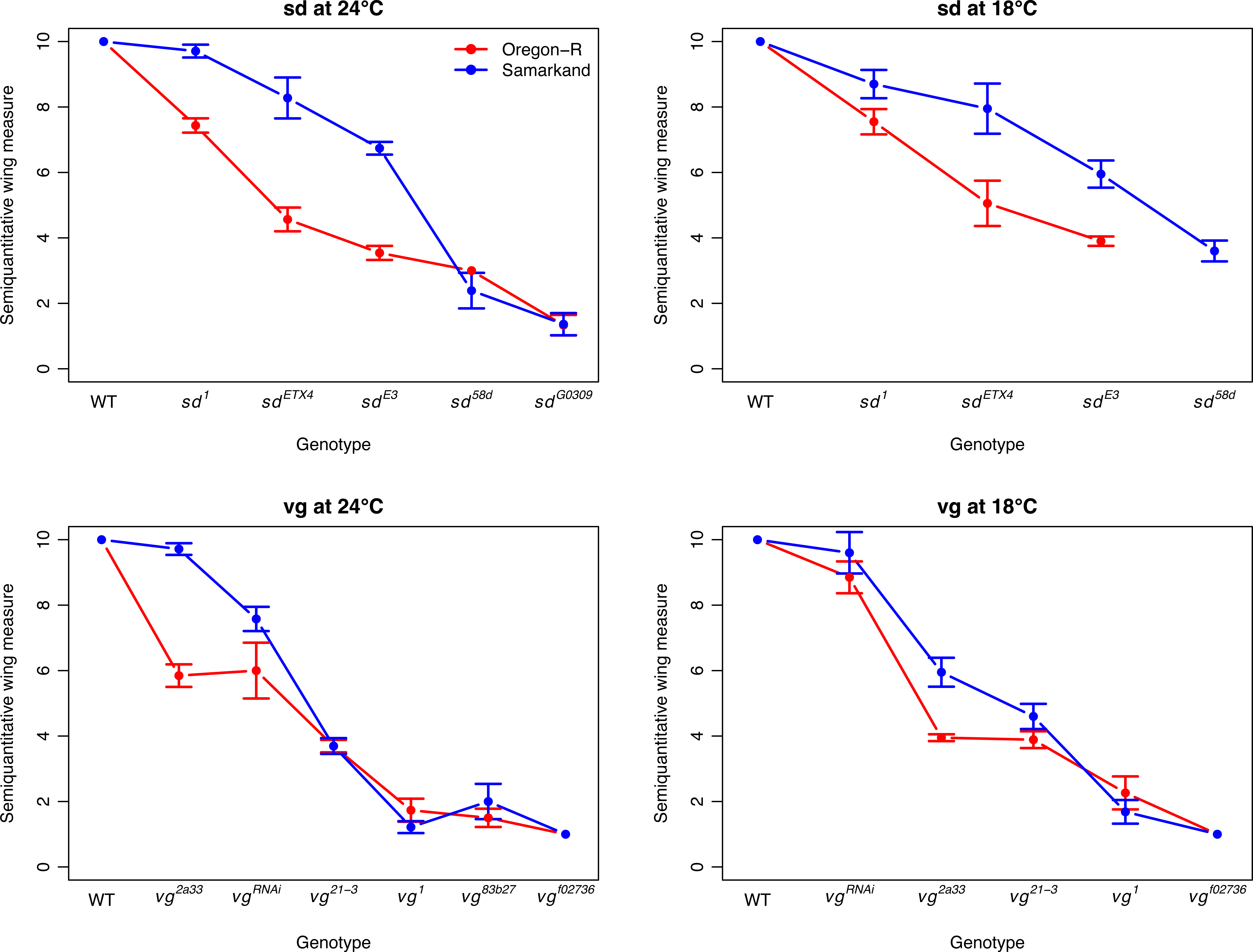
Genetic background and temperature effects of individual alleles of *sd* and *vg* without substantial re-ordering of the rank of allelic effects. The influences of genetic background and rearing temperature on the expressivity of allelic series in the *sd* and *vg* genes, using wing area as a measure of overall wing phenotype. "Stronger" mutations result in smaller wings. The *vgRNAi* represents a cross of the UAS_*vg*.RNAi to NP6333-GAL4 (both alleles introgressed into both genetic backgrounds). Error bars represent 95% confidence intervals.

**Supplemental Figure 3.**
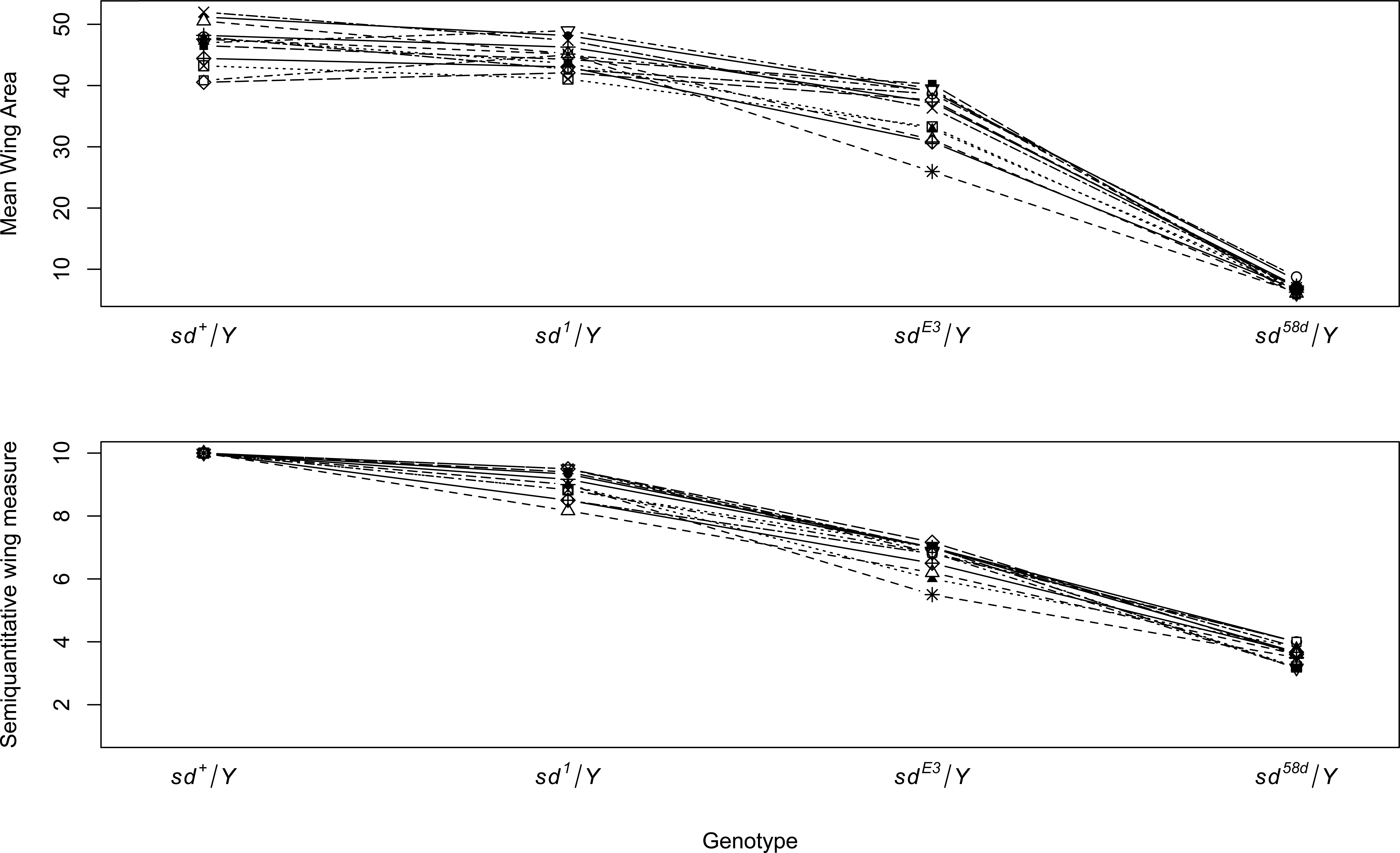
Ordering of allelic effects is maintained in crosses to 16 distinct wild type genotypes. Each line represents the male F1 offspring from crosses from males of one of 16 DGRP strains to females of SAM sd1, SAM sdE3, SAM sd58d (and corresponding 949 SAM wild type). Top panel is the measure of wing area, while the bottom panel uses the semi-quantitative measure of wing morphology. It is worth noting that the experimental design was by necessity different from the primary experiment (where alleles were introgressed and tested in homozygous genetic backgrounds). In this experiment, we are examining the effects of the sd alleles in hemizygous males, for the 16 genetic backgrounds heterozygous over the SAM genetic background. As such the among genetic background is far less (as recessive effects will generally not be expressed). Furthermore, we used the SAM background, which showed a generally weaker degree of phenotypic expressivity of mutations, which resulted in less severe phenotypic effects.

**Supplemental Figure 4.**
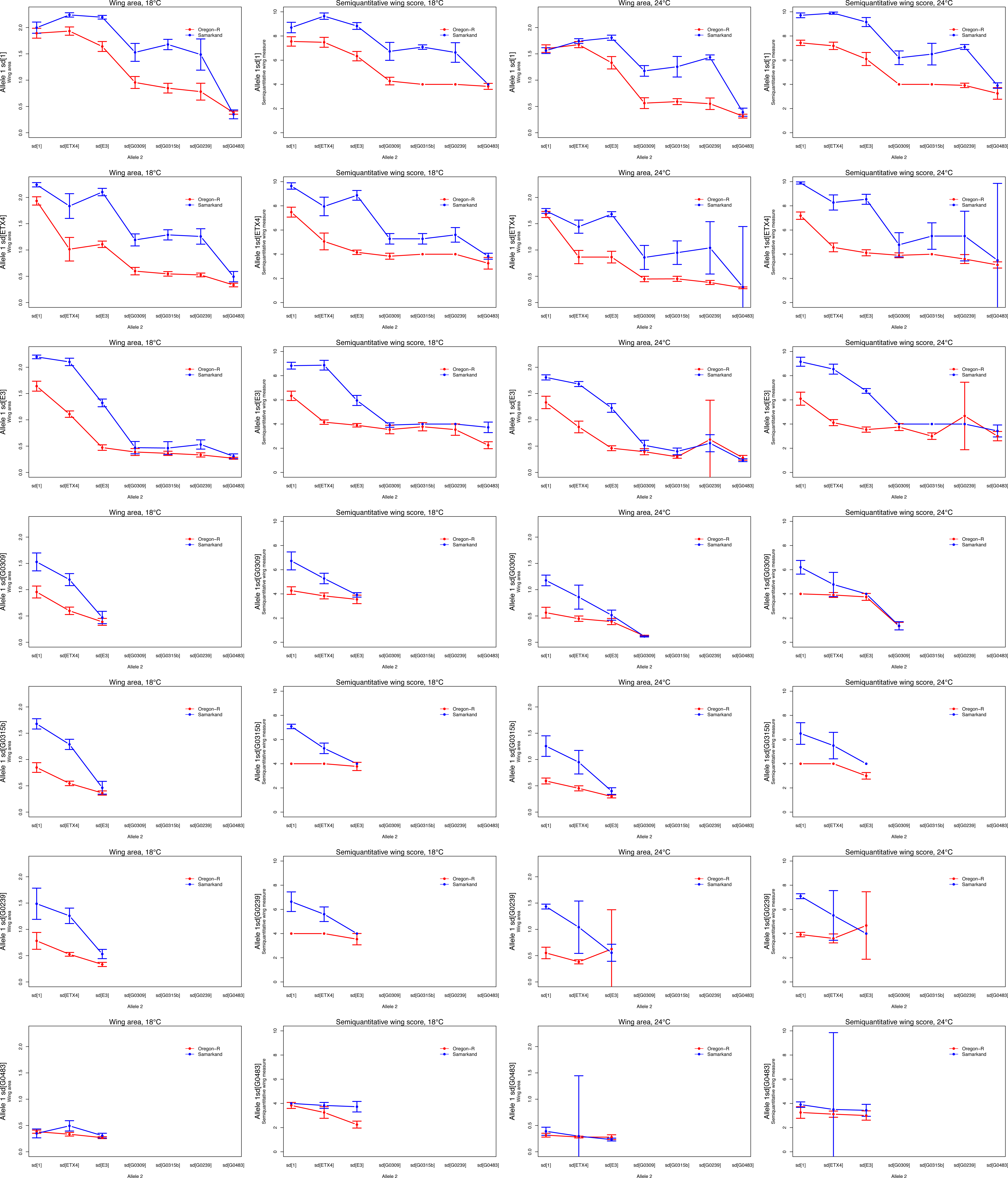
Full complementation plots for sd at both temperatures. Each panel depicts heterozygous flies carrying allele 1 (labeled on the y-axis) and allele 2 (x-axis). Left two panels: wing area and semiquantitative wing scores for flies reared at 18°C. Right 962 two panels: flies reared at 24°C. Error bars represent 95% confidenceintervals.

**Supplemental Figure 5.**
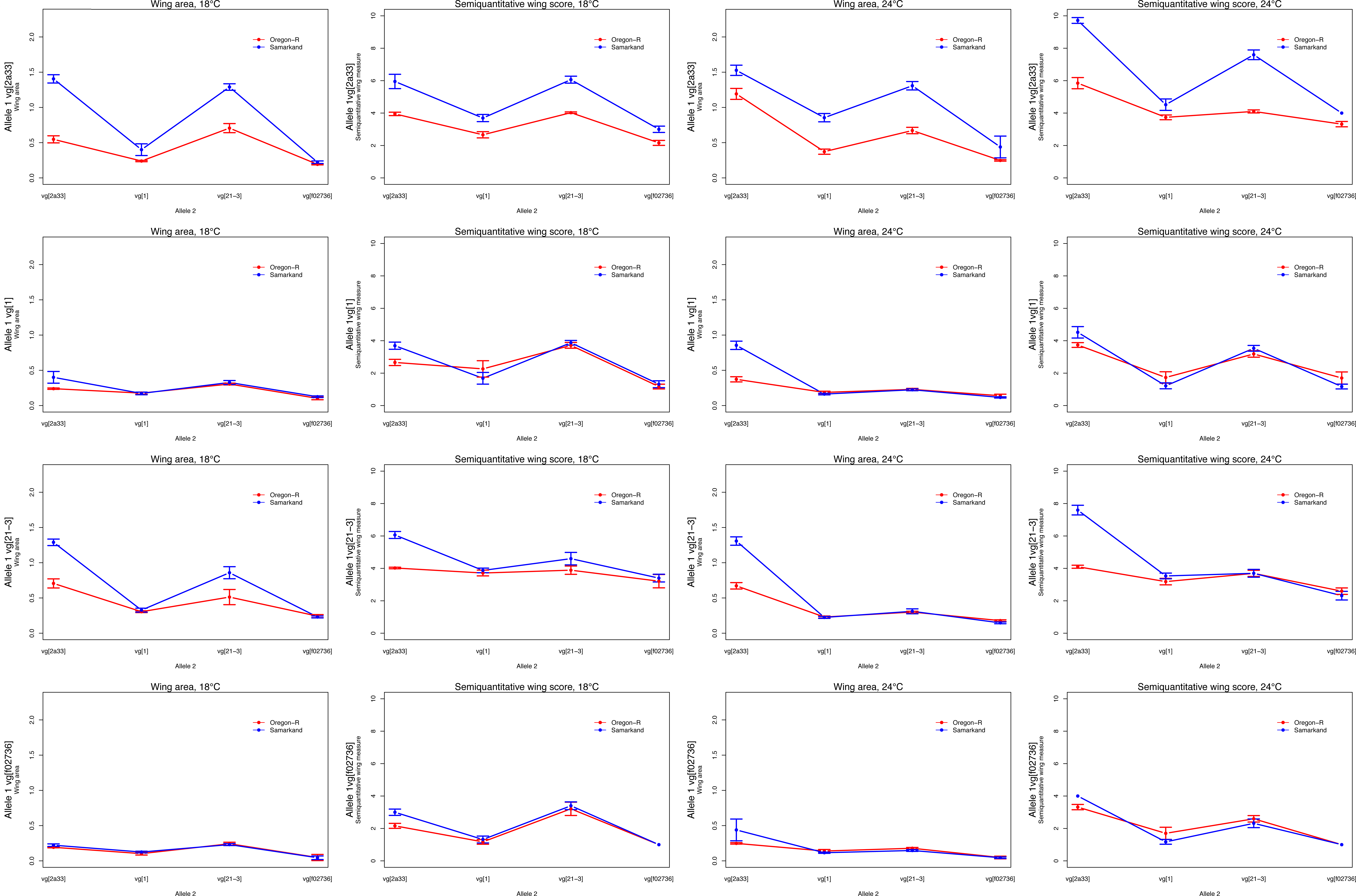
Full complementation plots for *vg* at both temperatures. Each panel depicts heterozygous flies carrying allele 1 (labeled on the y-axis) and allele 2 (x-axis).Left two panels: wing area and semiquantitative wing scores for flies reared at 18°C. Right two panels: flies reared at 24°C. Error bars represent 95% confidence intervals.

**Supplemental Figure 6.**
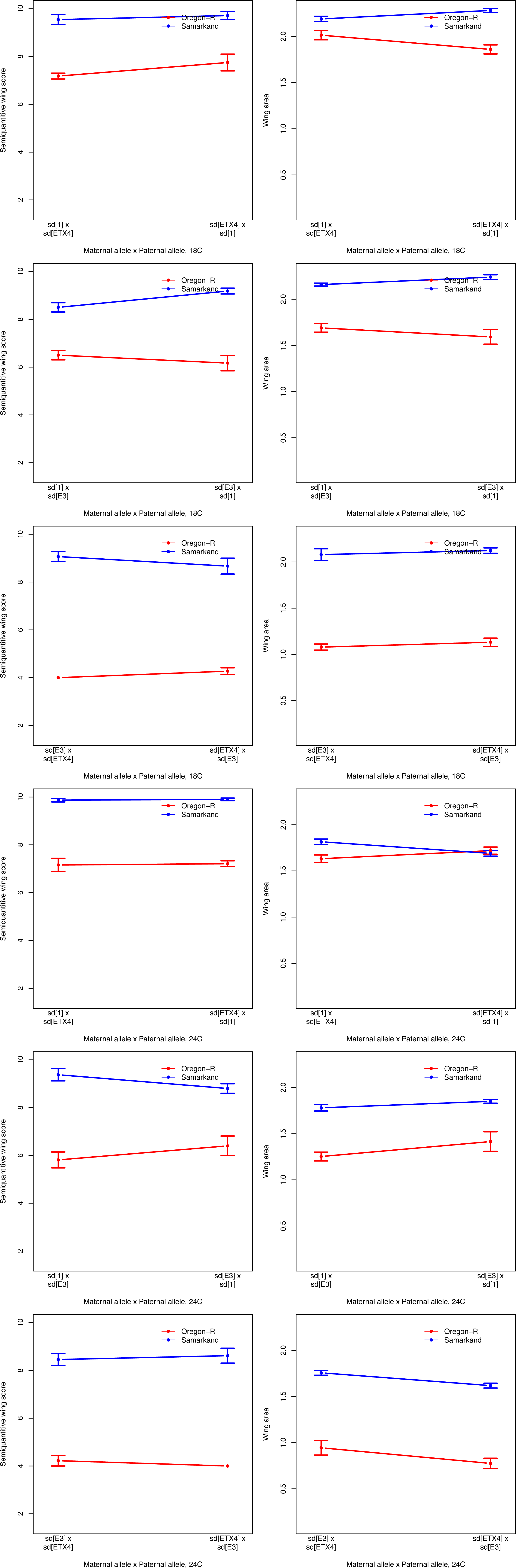
The effect of cross directionality on wing phenotypes was small; flies from reciprocal crosses between parents carrying different *sd* alleles have similar wing phenotypes. In this case, only female flies were measured,as *sd* is X-linked and male flies resulting from these crosses are hemizygous for just a single *sd* allele. Error bars represent 95% confidence intervals.

**Supplemental Figure 7.**
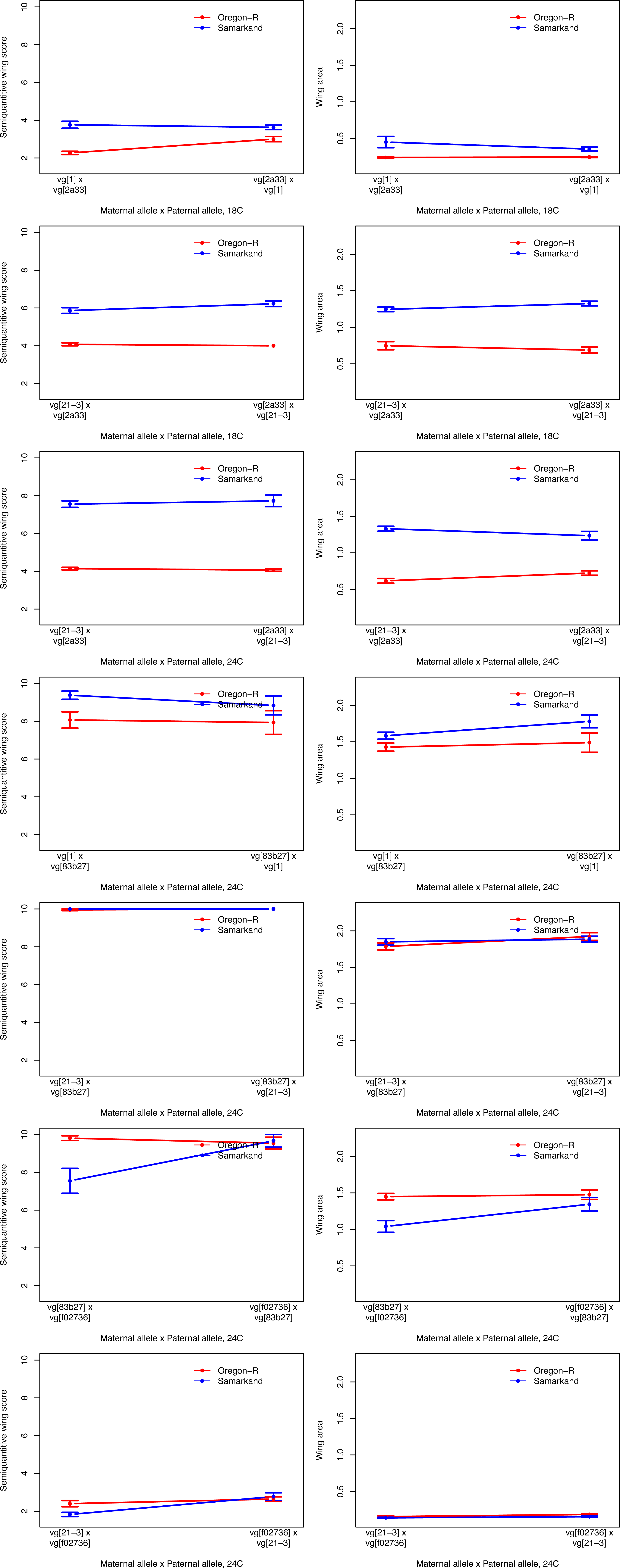
The effect of cross directionality on wing phenotypes was small; flies from reciprocal crosses between parents carrying different *vg* alleles have similar wing phenotypes. Error bars represent 95% confidence intervals.

**Supplemental Figure 8.**
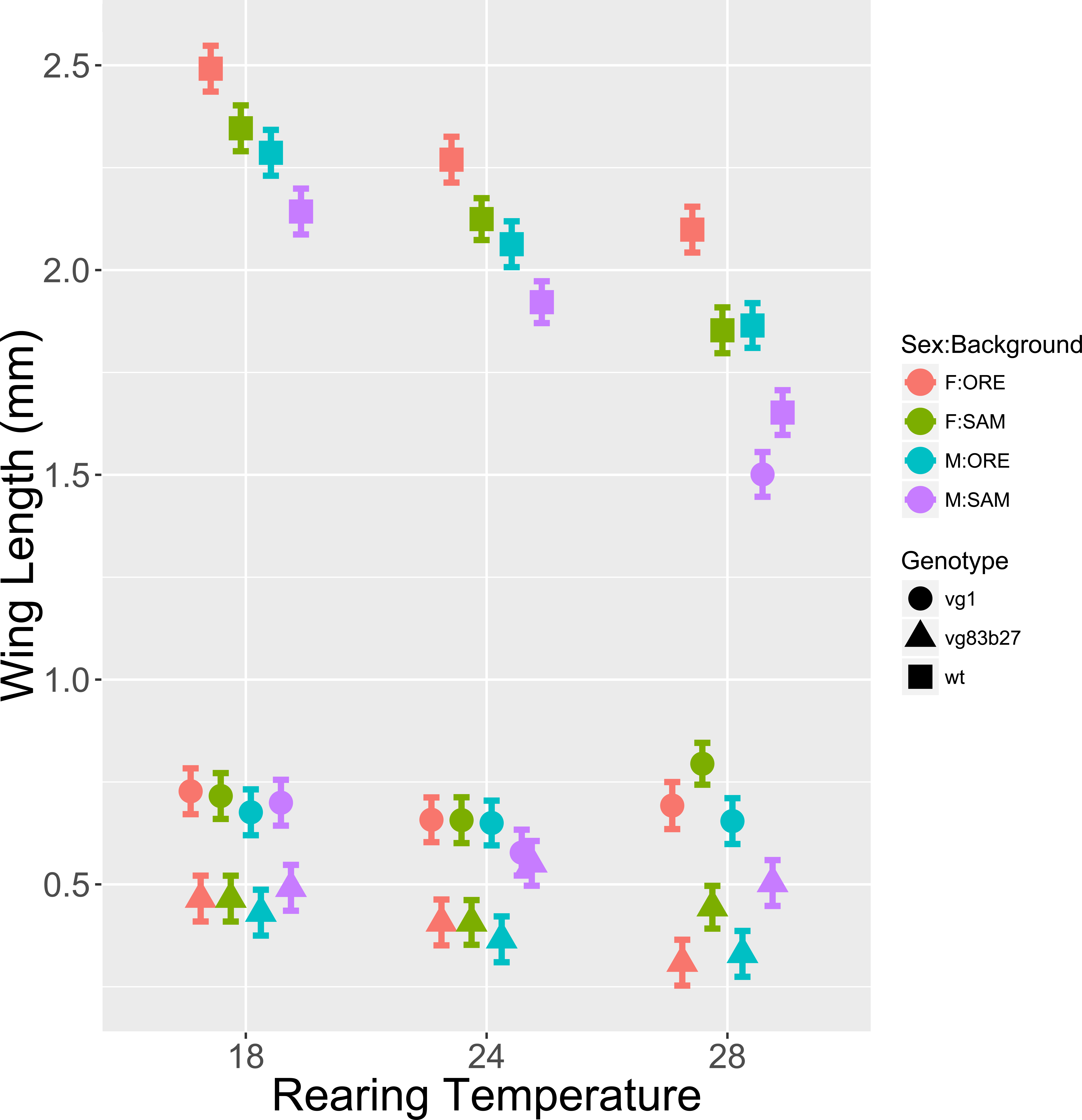
The phenotypic effects of the *vg1* allele are a function of rearing temperature, genetic background and sex. In particular note the phenotype of SAM *vg1* males at 28°C (purple circle). Error bars represent 95% CI.

**Supplemental Figure 9.**
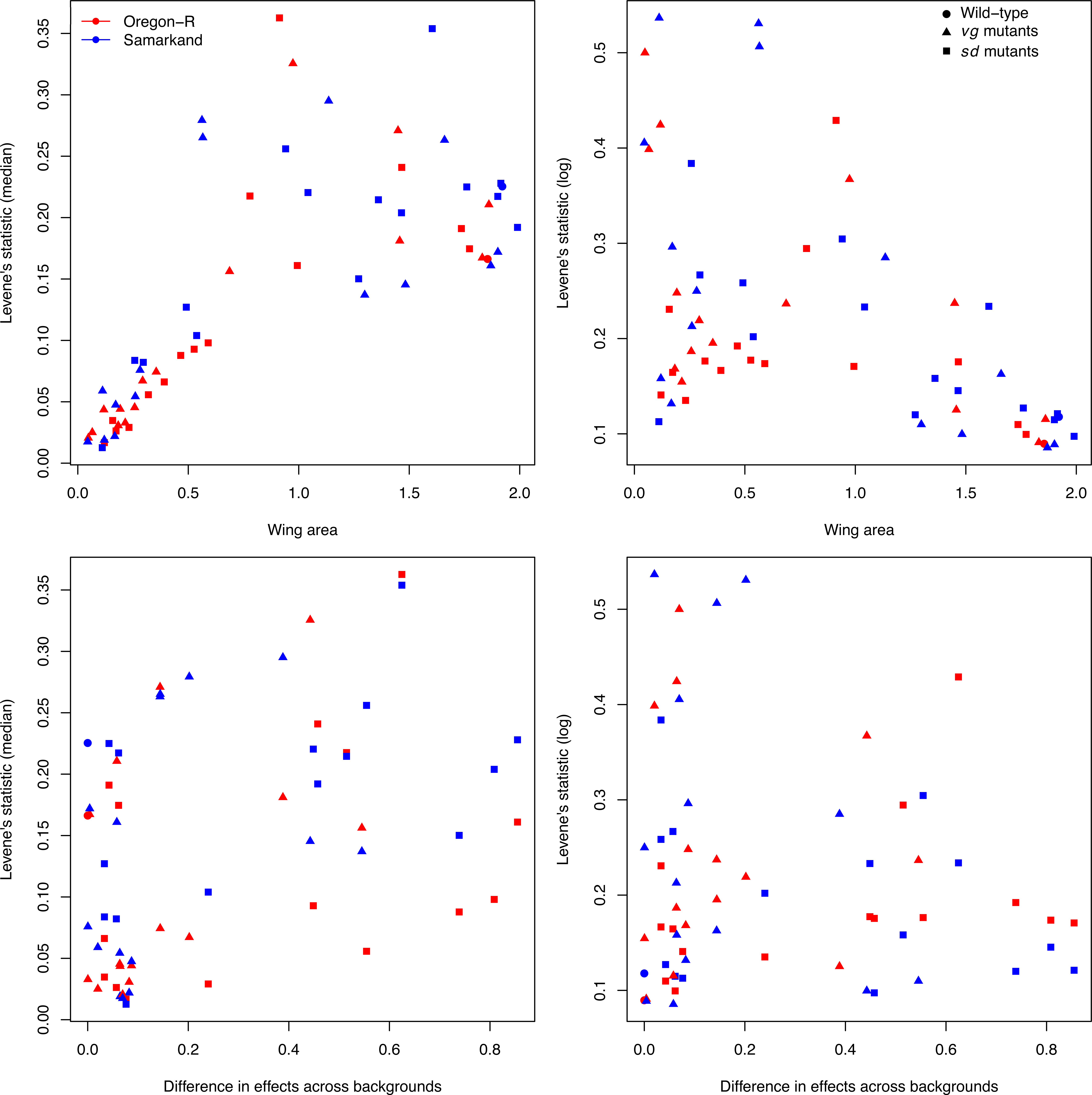
The relationship between among-individual, within-genotype/background variability and severity of mutational effects. There is a generally curvilinear relationship between the extent of mutational severity (as assessed by reduction in wing area) and intra-genotypic variability (A, B). As can be seen with the log transformed values of Levene’s statistic (B), genotypes with the most severe effects do increase variability, although with considerable among-genotype variation in the within-genotype variability. However, there is only a weak correlation between the background effects and intra-genotypic variability for the median form (C) of Levene’s statistic (Pearson r = 0.41, CI:0.18,0.6), and very weak for the log transformed measure (D) of Levene’s statistic (Pearson r = -0.09, CI: -0.33,0.16). We also observed little evidence for differences in the two wild type strains (ORE and SAM) in intrinsic variability across all genotypes. We used two forms of Levene’s statistic, one using raw deviations from the genotypic/background medians, and one using log transformed values to assess scaling effects for trait size.

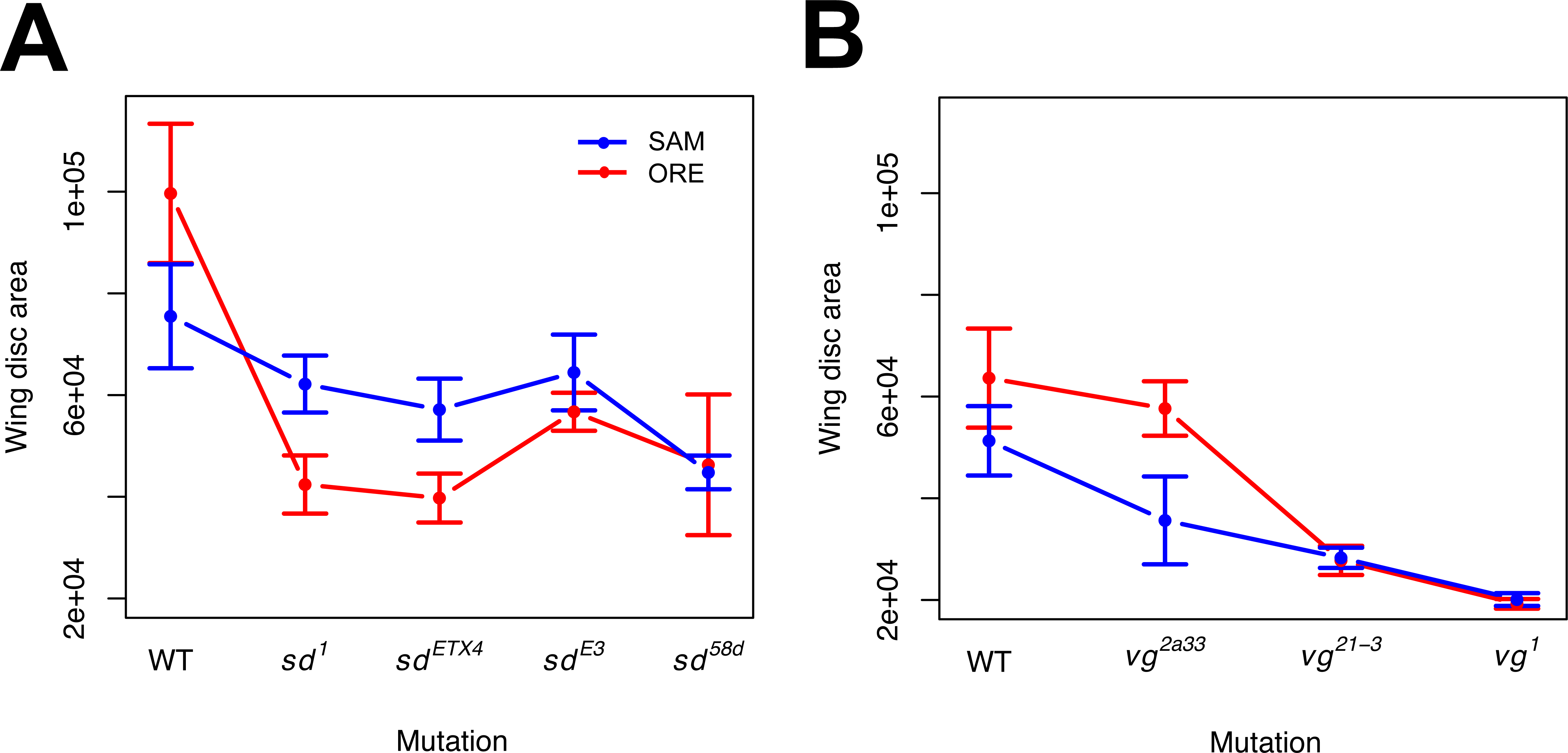

